# High-Speed, Cortex-Wide Volumetric Recording of Neuroactivity at Cellular Resolution using Light Beads Microscopy

**DOI:** 10.1101/2021.02.21.432164

**Authors:** Jeffrey Demas, Jason Manley, Frank Tejera, Hyewon Kim, Francisca Martínez Traub, Brandon Chen, Alipasha Vaziri

**Author notes:** Corresponding author: Alipasha Vaziri, Laboratory of Neurotechnology and Biophysics, The Rockefeller University, 1230 York Avenue, New York, NY 10065, USA.

## Abstract

Two-photon microscopy together with genetically encodable calcium indicators has emerged as a standard tool for high-resolution imaging of neuroactivity in scattering brain tissue. However, its various realizations have not overcome the inherent tradeoffs between speed and spatiotemporal sampling in a principled manner which would be necessary to enable, amongst other applications, mesoscale volumetric recording of neuroactivity at cellular resolution and speed compatible with resolving calcium transients. Here, we introduce Light Beads Microscopy (LBM), a scalable and spatiotemporally optimal acquisition approach limited only by fluorescence life-time, where a set of axially-separated and temporally-distinct foci record the entire axial imaging range near-simultaneously, enabling volumetric recording at 1.41 × 10^8^ voxels per second. Using LBM, we demonstrate mesoscopic and volumetric imaging at multiple scales in the mouse cortex, including cellular resolution recordings within ~3×5×0.5 mm^3^ volumes containing >200,000 neurons at ~5 Hz, recording of populations of ~1 million neurons within ~5.4×6×0.5 mm^3^ volumes at ~2Hz as well as higher-speed (9.6 Hz) sub-cellular resolution volumetric recordings. LBM provides an unprecedented opportunity for discovering the neurocomputations underlying cortex-wide encoding and processing of information in the mammalian brain.

## Introduction

Two-photon scanning fluorescence microscopy (2pM)^1–3^ combined with genetically-encoded calcium indicators (GECIs)^4–6^ as reporters of neuroactivity has emerged as the standard technique for imaging neuronal activity at depth and within scattering brain tissue. However, recent lines of evidence based on anatomical and functional observations suggest that complex brain functions emerge from highly parallel computation^7,8^ in which sensory information^9,10^ and behavioral parameters^11,12^ are mapped onto brain-wide neuronal populations^13–16^ beyond the scale of the small planar fields-of-view (FOVs) of conventional microscopes (< 0.5 mm). Maximizing volumetric FOVs toward brain-wide imaging requires both mesoscopic optical access as well as optimal spatiotemporal sampling, such that: (1) sample information is obtained at the fastest possible rate, (2) each voxel provides novel information about the sample at maximum signal-to-noise ratio (SNR), and (3) the microscope records only the minimum information necessary to resolve features of interest for the application (*e.g*. cell bodies) in order to devote remaining resources to imaging the largest possible volumes at calcium-imaging-compatible time scales.

Many platforms have demonstrated the requisite mesoscopic optical performance.^17–23^ However, in all of these systems spatiotemporal sampling remains suboptimal: the high repetition rate lasers employed lead to inevitable over-sampling in the lateral planes due to finite resonant scanner speed, and multiple-pulse-per-pixel binning to improve SNR at low pulse energies. Accordingly, performance is limited to at most multi-plane rather than volumetric performance, slow frame rates, and brain-heating-limited SNR, particularly when imaging at depth. As we have argued previously,^24,25^ single-pulse-per-voxel acquisition maximizes SNR per unit power delivered to the brain, and additionally, sampling at the minimum lateral density dictated by the application frees up temporal resources toward scaling volumetric FOV. Additionally, some systems have employed point-spread functions (PSF) that are intentionally extended along either the lateral^26^ or axial^27–29^ dimensions such that each voxel forms a projection of the volume along a given axis, reducing scanning requirements to two rather than three dimensions for volumetric operation. However, these approaches work best for sparsely-labeled samples, reducing applicability to imaging large networks, and also suffer from poor signal-to-background ratio when imaging deep in tissue.

Temporal multiplexing^30,31,32,19,23,25^ can be used to increase information throughput by scanning *N* copies of a single laser pulse which are delayed in time, and directed toward separate regions of the sample. As long as the relative delay between beams exceeds the fluorescence lifetime-limited decay time of the signal from the previous beam (~6-7 ns or ~140-160 MHz for GCaMP^31^), fluorescence at the detector can be re-assigned in order to reconstitute the FOV scanned by each beam, resulting in an *N*-fold increase in data rate beyond the inertial limits of the scanning system. However, multiplexed systems employing conventional high-repetition-rate lasers (typically ~80 MHz) are limited to at most a ~2-fold increase in data rate,^19,23,30–32^ in addition to suffering from oversampling inefficiencies as mentioned above. Lowering laser repetition rates to the few MHz regime can simultaneously increase the maximum possible degree of multiplexing *N* while improving sampling efficiency.^25^ However, many multiplexing platforms use feedforward approaches employing chains of beam splitters and dedicated optical paths for delaying and steering each beam, resulting in unfavorable scaling of system complexity with increasing *N*,^10,19,23,25,30,31^ and as a result can leave the majority of the inter pulse timing interval unexploited for maximizing information throughput.^25^ Recently, multi-pass temporal multiplexing schemes have been demonstrated using systems in which the beam is sent through a single set of components which tag the light with a specific delay and focal position in the sample corresponding to each pass through the system^33,34^. While using a multi-pass design it is in principle possible to achieve higher degrees of multiplicity at a reduced optical complexity, current realizations exhibit either fundamentally-limited potential for increase in multiplicity,^34^ or are inconsistent with both one-pulse-per-voxel excitation and axial multiplexing.^10,33^ Finally, spatial multiplexing – wherein copies of the input beam are not delayed, but instead, detection is spatially resolved – can also improve throughput,^35,36^ but demonstrations have been limited to shallow imaging depths due to scattering-induced spatial cross-talk between beams.

Here we demonstrate Light Beads Microscopy (LBM): a high-speed optical acquisition technique for simultaneously mesoscopic and volumetric 2pM. In LBM, the microscope scans a set of axially-separated and temporally-distinct foci (i.e. “beads”) as opposed to a single focus (Fig. 1a). The beads record information throughout the entire depth range of the sample (~500 μm) within the dwell time of a single pixel, thus LBM captures entire volumes within the time it takes to scan a single plane. Furthermore, by employing optimized spatial sampling, LBM can expand volumetric FOVs to mesoscopic scales while retaining GCaMP-compatible volume rates. Our light beads are formed by a cavity-based multiplexing approach called the Many-fold Axial Multiplexing Module (MAxiMuM). The distinct feature of the multiple-pass geometry of MAxiMuM is that it allows for scaling *N* to the limit posed by the fluorescence lifetime of GCaMP and the repetition rate of the laser, as well as controlling the relative power and position of each beam. It thereby provides 30-fold axial multiplexing and a voxel acquisition rate of 141 MHz with 16 μm plane-to-plane axial separation, conditions that are optimized for and compatible with sampling densely-labeled tissue volumes at fluorescence-lifetime-limited information rates, with one-pulse-per-voxel SNR-maximized excitation, while utilizing the entire inter pulse time interval.

**Fig. 1:**
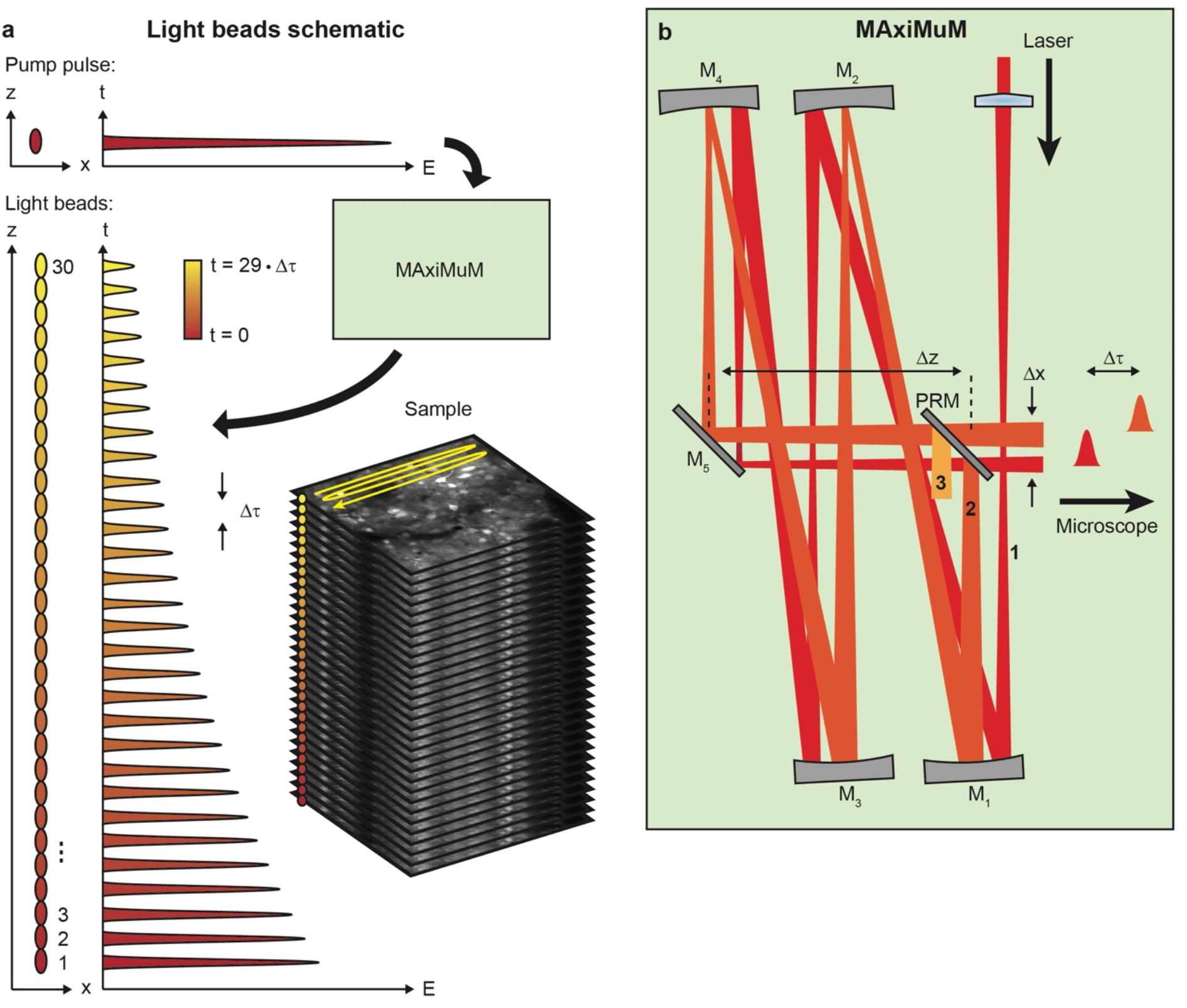
Light bead microscopy schematics. **a,** An ultra-fast pump pulse is split into 30 copies, which are delayed in time and sent to different depths in the sample, forming a column of light beads. Scanning the column samples the entire volume at the nominal frame-rate of the microscope. Each bead is temporally distinct, so fluorescence can be time-binned to decode the planes of the volume. **b,** Many-fold Axial Multiplexing Module (MAxiMuM) schematic. The red beam represents light entering the cavity, formed by four concave mirrors (M_1_-M_4_). A partially-transmissive mirror (PRM) re-injects most of the light back into the cavity. Beams accumulate an axial offset (*Δz*) and a temporal offset (*Δτ*) for each round trip, forming a column of light beads.

We have realized the design of our LBM on a mesoscopy platform that allows access to a lateral field of view (FOV) of ~6 × 6 mm^2^ at subcellular resolution (NA = 0.6),^18^ demonstrating volumetric and single-cell resolution recording from volumes of ~3 × 5 × 0.5 mm^3^, encompassing portions of the primary visual (VISp), primary somatosensory (SSp), posterior parietal (PTLp), and retrosplenial (RSP) areas of GCaMP6s-labeled^5,37^ mouse neocortex at ~5 Hz volume rate. Additionally, we highlight the versatility of LBM on this platform by recording in a variety of configurations ranging from moderately-sized FOVs (600 × 600 × 500 μm^3^) with voxel resolution capable of resolving subcellular features, to FOVs (5.4 × 6 × 0.5 mm^3^) encompassing both hemispheres of the mouse cortex and capturing the dynamics of populations exceeding ~800,000 neurons. We find that correlated activity of neurons in these experiments have characteristic lengths of » 1 mm, and stimulus- and behavior-tuned populations captured at mesoscale demonstrate richly varied responses at the single-trial level – including subpopulations with correlated trial-to-trial variations in their responses – underlining the need for such volumetric and mesoscopic calcium imaging techniques to understand real-time neurocomputations performed by inter-cortical circuitry.

## Results

### Light bead generation with MAxiMuM

The goal of LBM is to facilitate mesoscopic volumetric Ca^2+^ imaging compatible with the optimal spatiotemporal sampling conditions articulated above. To meet this goal, we constructed a stand-alone module called MAxiMuM capable of generating the requisite columns of multiplexed beams. Laser light is focused at the entrance into the MAxiMuM cavity and re-imaged by a series of concave mirrors (Fig. 1b, detailed schematic in Fig. S1). There is an intentional offset *Dz* between the nominal focus of the re-imaging mirrors and the input of the cavity, thus as the beam returns to the entrance, there is an axial shift in its focus. Before reaching the cavity entrance, the beam encounters a partially-reflective mirror which reflects the majority of light back into the cavity for another round trip, with a small lateral offset relative to first beam. The remaining fraction of the light couples out of the cavity and into the downstream microscope. Due to the focal offset, each beam exiting the cavity focuses to a shallower depth in the sample, with a relative decrease in optical power such that the power in the *i^th^* beam is given by *P_i_* = *T*(1 – *T*)^*i*^, where *T* is the transmission of the partially-reflective mirror.

It is well known in 2pM that maintaining SNR at depth requires an exponential increase in laser power owing to loss of ballistic photons due to tissue scattering. It follows that the transmission of the cavity, *T*, can be adjusted such that the relative increase in power for each sequential beam matches the increase required to offset additional tissue scattering due to the axial separation between adjacent planes, *δz*, and thus achieve constant SNR for all multiplexed beams. This condition is given by:

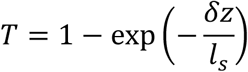

where *l_s_* is the scattering mean free path of brain tissue (~200 μm for a laser wavelength of 960 nm^38^), and the relationship between the axial separation of beams exiting the cavity (*Dz*) versus in the sample (*δz*) is given by *Dz* = *M^2^δz*, where *M* is the lateral magnification of the microscope. Equation (1) provides a design rule for achieving a given axial sampling density with the LBM approach (see Supplementary Note 1).

We designed our MAxiMuM module based on, and to be integrated into, an existing mesoscope (NA = 0.6, FOV = 6 mm, Fig. S2) and characterized each light bead in sample space (Fig. S3, Supplementary Note 2) to ensure desired temporal and spatial characteristics. Through these calibrations, we confirmed fluorescence-lifetime-limited bead-to-bead delays (6.7 ns) and minimal cross-talk between channels while the axial location at which our beads focused increased linearly with bead number for our 30 light beads over a total axial range of ~500 μm. We further confirmed a lateral and axial full-width-at-half-maximum localization diameter of our light beads of ~1 μm and ~13 μm, respectively; sufficient for cellular-resolution imaging of densely labeled samples.

### Optimization of spatiotemporal sampling efficiency

In order to maximize spatiotemporal sampling efficiency and record from the largest possible FOV, only the minimum information necessary to faithfully extract features of interest – in our case, neuronal cell bodies – should be recorded. Accordingly, we set out to determine the minimal spatial sampling requirements in order to resolve the GCaMP transients of neocortical mouse neurons. We recorded several high-resolution single plane data stacks and fed them into our analysis pipeline (Fig. S4, see Supplementary Note 3 for details), comparing the extracted footprints and time series to hand-segmented functional ground truths for each data set. Using F-score (defined as the harmonic mean of the true and false positive rates) as a metric for extraction fidelity, we evaluated how performance deteriorates as a function of sampling sparsity by removing pixels from the lateral image stacks. Consistent with our previous results,^24,25^ we found that the F-score only significantly declines for lateral spatial sampling > 5 μm (Fig. S5). Thus, to maximize the imaging volume while maintaining a high signal extraction fidelity, we found an optimum sampling of ~5 μm in the lateral plane.

### Multi-regional and multi-sensory imaging of activity from >200,000 neurons in mouse cortex

We validated LBM by performing *in vivo* imaging in the neocortex of awake and behaving mice transgenically expressing GCaMP6s.^5,37^ Using our optimized spatial sampling strategy in a volume of ~3 × 5 × 0.5 mm^3^ we could maintain a ~5 Hz frame rate compatible with resolving GCaMP transients. We oriented the FOV in order to encompass as many distinct regions as possible within a single cortical hemisphere including SSp and PTLp, as well as RSP and VISp (Fig. 2a). In order to evoke activity of neurons distributed through the many functional regions within the FOV, we developed a dual sensory stimulus paradigm including whisker and visual stimulation. During whisker trials, we perturbed the majority of the whiskers contralateral to the hemisphere being imaged. Each visual trial consisted of a high-contrast drifting grating with orientation following a sequence of horizontal, 45°, vertical, 135° which was repeated *ad nauseum* throughout the recording. The overall stimulus paradigm consisted of a whisker trial, a visual trial, followed by a simultaneous whisker and visual trial, with 5 s intervals between each trial. Additionally, our treadmill was equipped with velocity tracking and video recording of the head-fixed animals in order to capture and track their spontaneous behaviors, such as paw and torso movements (Movie S1). Typical recordings in this modality ranged from 9–30 minute duration, yielding populations of 159 – 460,000 neurons (12 recordings from N = 6 mice, see Fig. S6a).

**Fig. 2:**
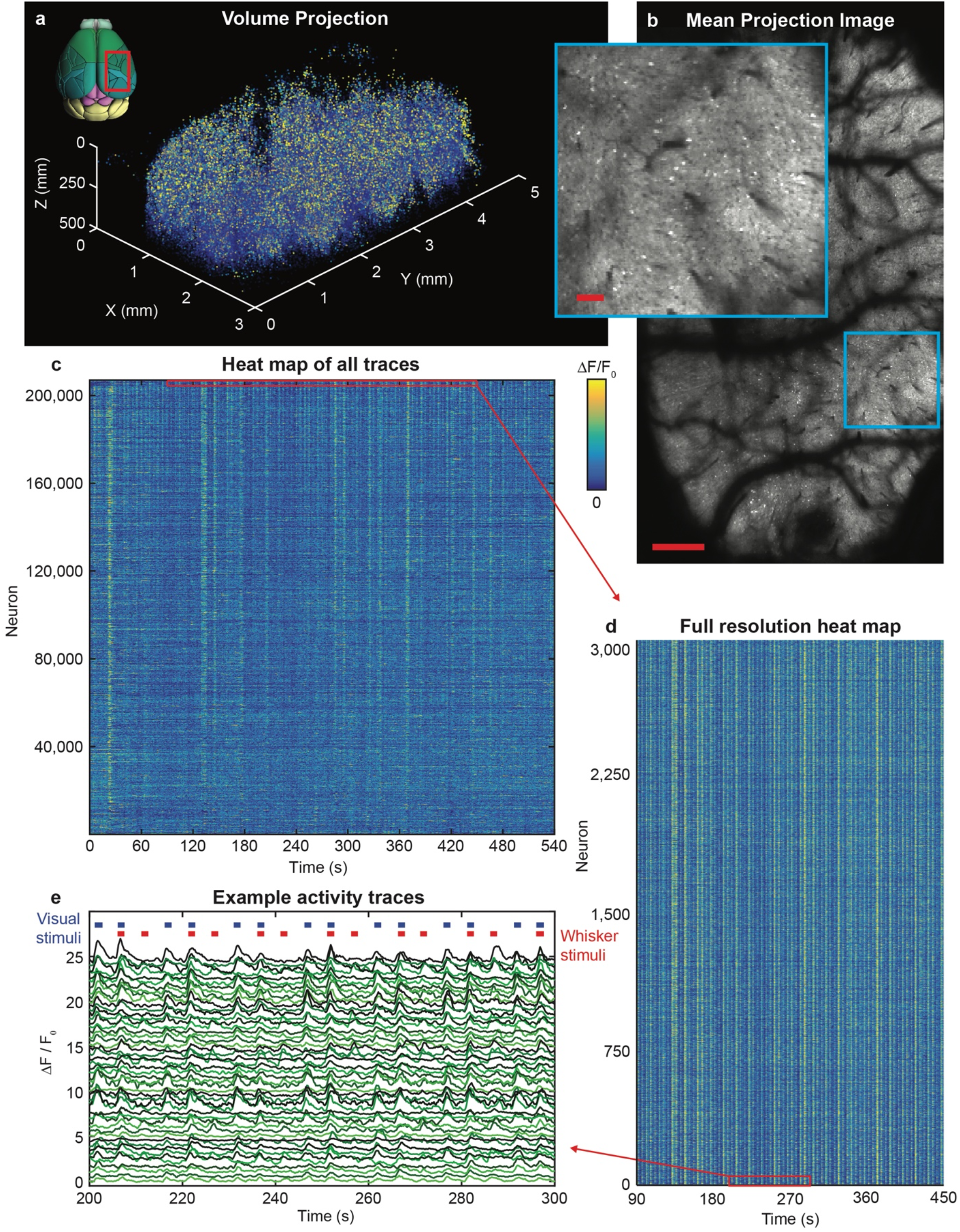
Recording of 207,030 neurons within a ~3 × 5 × 0.5 mm^3^ volume in a GCaMP6s-expressing mouse at 4.7 Hz rate. **a,** 3D rendering of extracted neuron spatial coordinates and maximum projected activity for a 9-minute recording. Transverse brain image reproduced from Ref. 52. See also Movie S2. **b,** Mean projection image from the recording in a at 344 μm depth. Scale bar: 250 μm. Inset: zoomed in region of **b**, scale bar: 100 μm. See also Movie S3. **c,** Heat map of all 207,030 neurons extracted from the recording in **a** normalized to maximum *ΔF/F_0_*. **d,** Subset of 3,000 traces from **c** shown at full resolution. **e,** Subset of 50 traces from **d** with whisker and visual stimuli denoted by red and blue markers, respectively. Offset: 0.5 *ΔF/F_0_*.

Figure 2 shows data from a representative 9-minute recording of 207,030 neurons distributed across a ~500 μm axial range within the multiple cortical regions mentioned above (volume rendering in Fig. 2a and Movie S2). As expected, (Fig. S5), and consistent with the spatiotemporal characteristics of the light beads generated by MAxiMuM (Fig. S3), individual neurons could indeed be faithfully resolved (Fig. 2b, Movie S3), allowing for high fidelity extraction of their time-series (Figs. 2c–2e). Due to the sheer amount of data, it is difficult to display the time-series of all 207,030 neurons at full resolution simultaneously. Figure 2c shows an overview heatmap of all the neurons (see Movie S4), while Fig. 2d shows a detailed heatmap of a subset of 3,000 neurons exhibiting periodic stimulus-evoked activity. Traces for a further subset of 50 neurons are shown in more detail in Fig. 2e with visual and whisker trials denoted by blue and red marks, respectively. The time-scale of the observed calcium transients are consistent with the characteristic response time of GCaMP6s (Fig. S6) and show correlation with both visual and whisker stimuli.

To further investigate the tuning of neurons to external variables, we computed the distribution of correlations between the time-series of each neuron with lag-corrected, GCaMP-response-kernel-convolved time vectors associated with the presentation of each of the stimuli, *i.e*. the whisker trials, the visual trials, as well as correlation with spontaneous, uninstructed animal behaviors including movements of the fore or hindlimbs (Figs. S7a–7c). We found subpopulations of neurons spanning many regions of the cortex (Fig. 3a–3e) that were highly tuned to each stimulus (*R* > *3σ*, where *σ* is the standard deviation of a normal distribution fit to the distribution of correlations between neuronal time series and a time-shuffled stimulus vector, see Methods for details), numbering 34,468 whisker-tuned neurons, 24,299 visual-tuned neurons, and 64,810 neurons tuned to uninstructed animal behaviors.

**Fig. 3:**
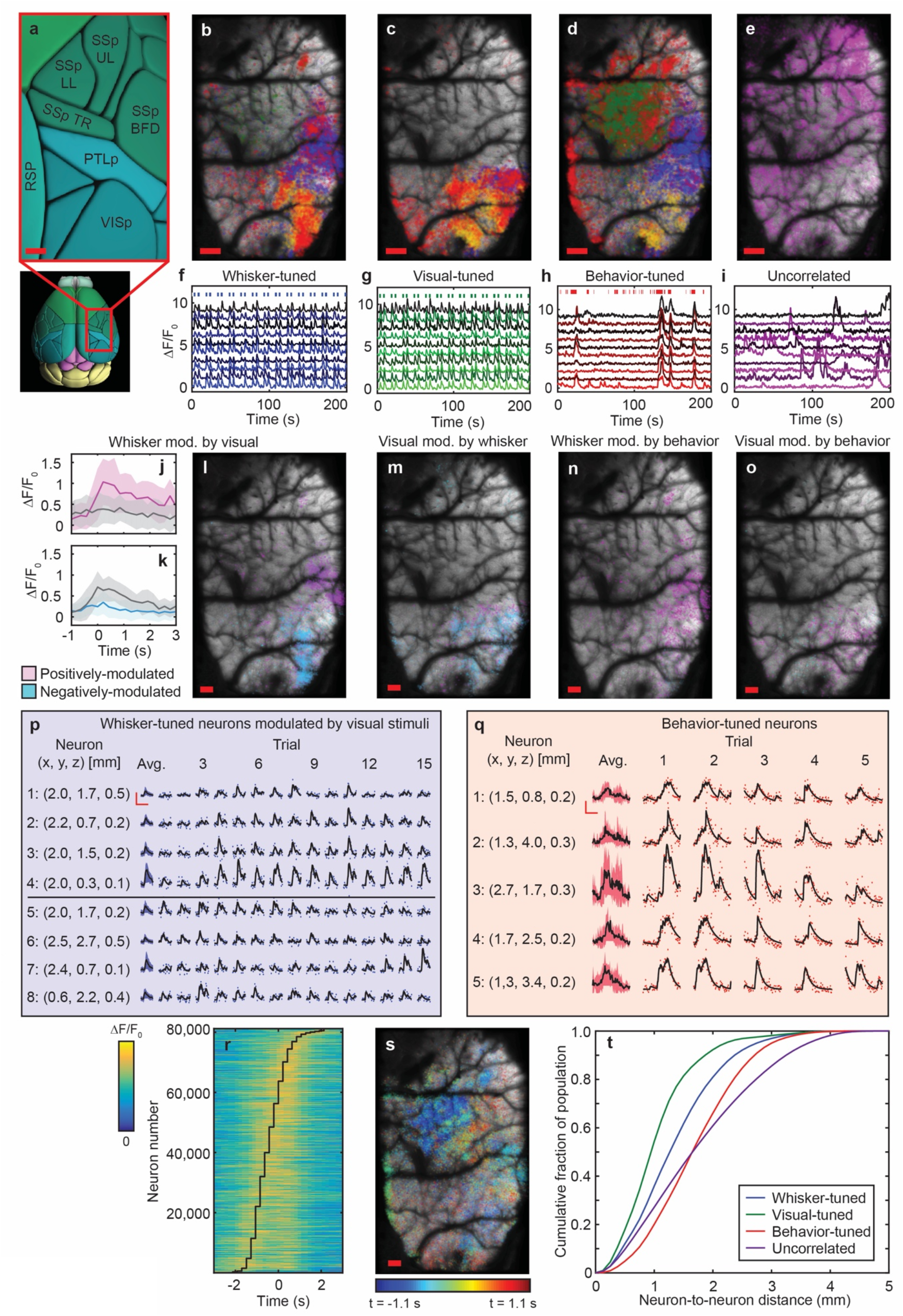
Analysis of the activity of stimulus-tuned and behavior-correlated neurons in a single hemisphere recording. **a,** Brain regions covered by the recording shown in Fig. 2, reproduced from Ref. 52. Scale bar: 250 μm. **b–e,** Transverse spatial distributions of neurons tuned to a single stimulus condition. The correlation matrix (see Fig. S7) for all tuned neurons was hierarchically clustered and sorted by preferred stimulus generating 4 clusters represented by blue, green, yellow, and red coloring, respectively, throughout the figures. Each map corresponds to whisker stimuli, visual stimuli, or behaviors respectively, with **e** corresponding to a population of neurons not correlated with any stimuli. Scale bars: 250 μm. **f–i,** Example neuronal traces from populations tuned to whisker stimuli, visual stimuli, spontaneous behaviors, or uncorrelated, respectively. Occurrence of stimuli denoted by markers in appropriate cases. Offset: 1.0 *ΔF/F_0_*. **j,k,** Trial-averaged activity of example whisker-tuned neurons with (magenta in **j**, cyan in **k**) and without (gray) the presence of a simultaneous visual trial. In **j**, the activity of a whisker-tuned neuron is positively-modulated – i.e. activity increases – when visual stimuli are simultaneously presented; in **k**, a different whisker-tuned neuron is negatively-modulated. Solid lines denote mean of all trials, shaded regions denote 1 standard deviation from mean. **l,** Lateral spatial distributions of the magenta and cyan populations in **j** and **k**. Scale bar: 250 μm. **m,** lateral spatial distributions of visually-tuned neurons positively- and negatively-modulated by simultaneous whisker stimuli. **n,** lateral spatial distributions of whisker-tuned neurons positively- and negatively-modulated by simultaneous animal behaviors. **o,** lateral spatial distributions of visually-tuned neurons positively- and negatively-modulated by simultaneous animal behaviors. **p,** single trial activity for example neurons tuned to whisker stimuli for trials with simultaneous visual stimuli. The left-most transient shows the trial averaged activity with the shaded portion denoting 1 standard deviation from the mean. The raw data for each example transient in trials 1–15 is shown by markers, black lines denote the deconvolved response; coordinates for each neuron are listed in units of mm for each neuron; horizontal and vertical scale bars: 1.0 *ΔF/F_0_*, 5 s. **q,** single trial activity for example neurons tuned to uninstructed behaviors of the animal; left-most transient shows the trial averaged activity with the shaded portion denoting 1 standard deviation from the mean; raw data for each example transient in trials 1–5 is shown by markers, black lines denote the deconvolved response; coordinates for each neuron are listed in units of mm for each neuron; horizontal and vertical scale bars: 1.0 *ΔF/F_0_*, 5 s. **r,** Heat map of trial-averaged activity of behavior-tuned neurons with relative lag denoted by the overlaid black line. Heat map is normalized to maximum *ΔF/F_0_* for each neuron and trial averaging includes the top 5 highest velocity behavior trials. **s,** Lateral spatial distribution of behavior-tuned neurons color-coded by relative lag. Scale bar: 250 μm. See also Movie S5. **t,** Cumulative fraction of populations tuned to a given condition (whisker stimulus, visual stimulus, spontaneous behavior, uncorrelated) with significant mutual correlation (*R* > *3σ*) captured within a given neuron-to-neuron separation.

We performed hierarchical clustering on the mutual correlation matrix of the population of all 123,577 stimulus-tuned neurons (Fig. S7d) and found 4 distinct clusters determined by ensuring similar node-to-stem distances for each resulting branch in the dendrogram. We subsequently mapped these clusters, and the neurons within them, back to their anatomical locations in the brain. Figures 3b–3d show the cortex-wide lateral spatial distributions of neurons associated with a given condition (whisker stimulation, visual stimulation, and uninstructed spontaneous behavior respectively), color-coded by their respective clusters, while Fig. 3e shows the lateral distribution of 13,259 neurons uncorrelated with any stimulus condition (|*R*| < *σ*). For the majority of the stimulus conditions, a distribution of correspondingly tuned neurons across multiple regions of the cortex (Fig. 3a) can be observed. For each condition, as well as for the uncorrelated population, we can faithfully extract the transients of single cells as examples shown in Figs. 3f–3i respectively demonstrate.

Cluster 1 (blue) was located primarily in the barrel field (SSp-BFD) and PTLp (Fig. 3a), and accordingly was highly represented in the whisker-tuned population (Fig. 3b). This cluster was also highly represented in the population correlated with spontaneous behaviors (Fig. 3d), inferring mixed response of the neurons in this cluster to both stimuli. Cluster 2 (green) was only represented in behavior-tuned neurons (Fig. 3d) and primarily located in specialized regions of the SSp related to sensation in the lower limbs, upper limbs, and torso of the animal (SSp-LL, SSp-UL, and SSp-TR, respectively), as well as PTLp (Fig. 3a). Cluster 3 (yellow) was located in VISp and PTLp, and represented neurons correlated with all stimulus conditions. The presence of visually-associated neurons in the whisker-tuned population may have been due to the fact that animal could see the motion of the brush stimulating the whiskers during stimulus presentation. The final cluster 4 (red) was distributed across multiple regions, including SSp, VISp, PTLp, and a dense population within RSP, which is thought to be associated with spatial memory encoding.^39^ This subset located within RSP was found to be primarily tuned to uninstructed spontaneous behaviors (Fig. 3d). The spatial clustering analysis suggests that, while some of these functional clusters overlap with distinct anatomical regions of the brain, neurons in these regions can also jointly represent multiple stimulus conditions or may have stimulus-evoked activity that is further modulated by the presence of additional stimuli.

To probe mixed representation further, we analyzed the trial-averaged activity of stimulus-tuned neurons. First, we considered the difference in trial-averaged activity between whisker-tuned neurons for trials where only the whisker stimulus was present compared to those trials where both whisker and visual stimuli occurred simultaneously (Figs. 3j–3l). Example trial-averaged transients for both a positively and negatively modulated neuron are shown in Figs. 3j and 3k respectively. The lateral spatial locations of significantly (*p* < 0.05, determined by t-test) upward and downward modulated neurons are shown in Fig. 3l. There were roughly similar numbers of whisker-tuned neurons positively (3,703 neurons) and negatively modulated (4,166 neurons). However, there was a clear distinction between the anatomical location of the two populations, with positively-modulated neurons located primarily in SSp-BFD and negatively-modulated neurons located in VISp. Figure 3m shows a map of visually-tuned neurons with activity significantly modulated by coincident presentation of whisker stimuli. Visually-tuned neurons were primarily negatively-modulated by the presence the of whisker stimuli, and located within VISp. Figures 3n and 3o show the population of whisker- and visually-tuned neurons significantly modulated by coincident uninstructed spontaneous behaviors of the animal. In both cases, the majority of whisker- and visually-tuned neurons are positively-modulated by spontaneous behaviors.

Additionally, at the single trial level, stimulus- and behavior-tuned populations show significant neuron-to-neuron and trial-to-trial variation. Figure 3p shows example traces from eight neurons tuned to whisker-stimuli in the presence of coincident visual-stimuli. In all cases, the response of a given neuron varies significantly from trial to trial. In some instances, neurons anatomically separated by >1 mm exhibit variations in activity across trials that are correlated (neurons 1–4), while in other instances the variations in trial-to-trial activity do not covary with the above group nor one another (neurons 5–8). Figure 3q shows example single trial responses of neurons tuned to uninstructed behaviors. Rather than just variations in the magnitude of single calcium transients, the response of these neurons also show variability in the total number of transients as well as the onset time of their responses across trials. We quantified this variability in onset time by calculating the lag of the onset time of the activity of each behavior-tuned neuron with respect to the onset of the behavior and found a ~1.7 s delay between the timing of the peak activity of the earliest and latest neurons. Fig. 3r shows the heatmap of the behavior-tuned population sorted by preferred lag and trial-averaged over the top 5 highest velocity movements while Figure 3s shows the lateral positions of these behaviors-tuned neurons color-coded by relative onset time (Movie S5). The earliest-responding neurons are primarily located in the SSp-TR, SSp-LL, and SSp-UL regions of the brain, while neurons in regions farther from these sensory areas, including RSP, PTLp, SSp-BFD, and VISp, respond significantly later, in keeping with previous results in the literature.^11^

The spatial and temporal structure of neurons captured in these recordings highlight the need for mesoscale and volumetric recording capabilities. Stimulus-tuned, behavior-tuned, and uncorrelated neurons in the recording shown in Figs. 2 and 3 exhibit correlated activity that spans neuron-to-neuron separations of 2–4 mm (Fig. 3t), and thus require mesoscale recording to capture the dynamics of the entire population. Additionally, stimulus-tuned neurons exhibit a trial-by-trial variability of response that appear to covary across the population, despite being separated by pair-wise distances on mesoscopic scales. Such trial-to-trial covariations of neuronal responses, also referred to as “noise correlations” have been suggested to represent distributed, higher dimensional encoded information underlying the interaction of external stimuli and behavioral states with internal states, or to represent information related to uncontrolled aspects of the stimulus presentation or untracked behavioral states.^11,40^ Due to the inherent nature of noise correlations which vary on a trial-by-trial basis, any trial averaging will fundamentally prevent their detection. Thus, while sequential single plane and tiled FOV recordings could potentially capture the same population of cells shown here, the trial-to-trial variability of their responses recorded by our method would be inherently lost in such an approach. Furthermore, since these noise correlations are thought to encode information about brain state, uninstructed behaviors, or unintended stimuli, it follows that disambiguating populations of neurons with correlated variation (neurons 1–4, Fig. 3p) from those with uncorrelated trial-to-trial activity (neurons 5–8, Fig. 3p), as well as the robustness of any information carried by projections of the full population dynamics, would improve as the number of neurons recorded increases in a similar manner to traditional stimulus encoding in primary sensory areas.^9,40^

### Re-configurable multi-scale imaging with LBM

LBM maintains the capability to navigate tradeoffs between lateral voxel sampling, FOV, and imaging speed to suit numerous applications. By decreasing the stroke of the scan mirrors, one can increase sampling density at the cost of FOV, and by adding more lateral scanning regions, one can increase FOV at the cost of volume rate. For example, Figs. 4a–4c show mean projection images and example neuronal traces from a volume of ~600 × 600 × 500 μm^3^ in the PTLp of a jGCaMP7f^41^ expressing mouse (see also Movie S6) at ~10 Hz volume rate and 1 μm lateral voxel sampling, sufficient for resolving sub-cellular features such as the fine dendrites of active cells. Relaxing voxel sampling to an intermediate ~3 μm lateral sampling (Figs. 4d–4g, Movies S7 and S7) allows for increasing the recording volume to ~2.0 × 2.0 × 0.5 mm^3^ at a volume acquisition rate of 6.5 Hz allowing to capture the activity of a population of ~70,000 neurons in GCaMP6s transgenically-labeled mice at imaging resolutions higher than what would be needed to resolve typical mouse cortical neurons.

**Fig. 4:**
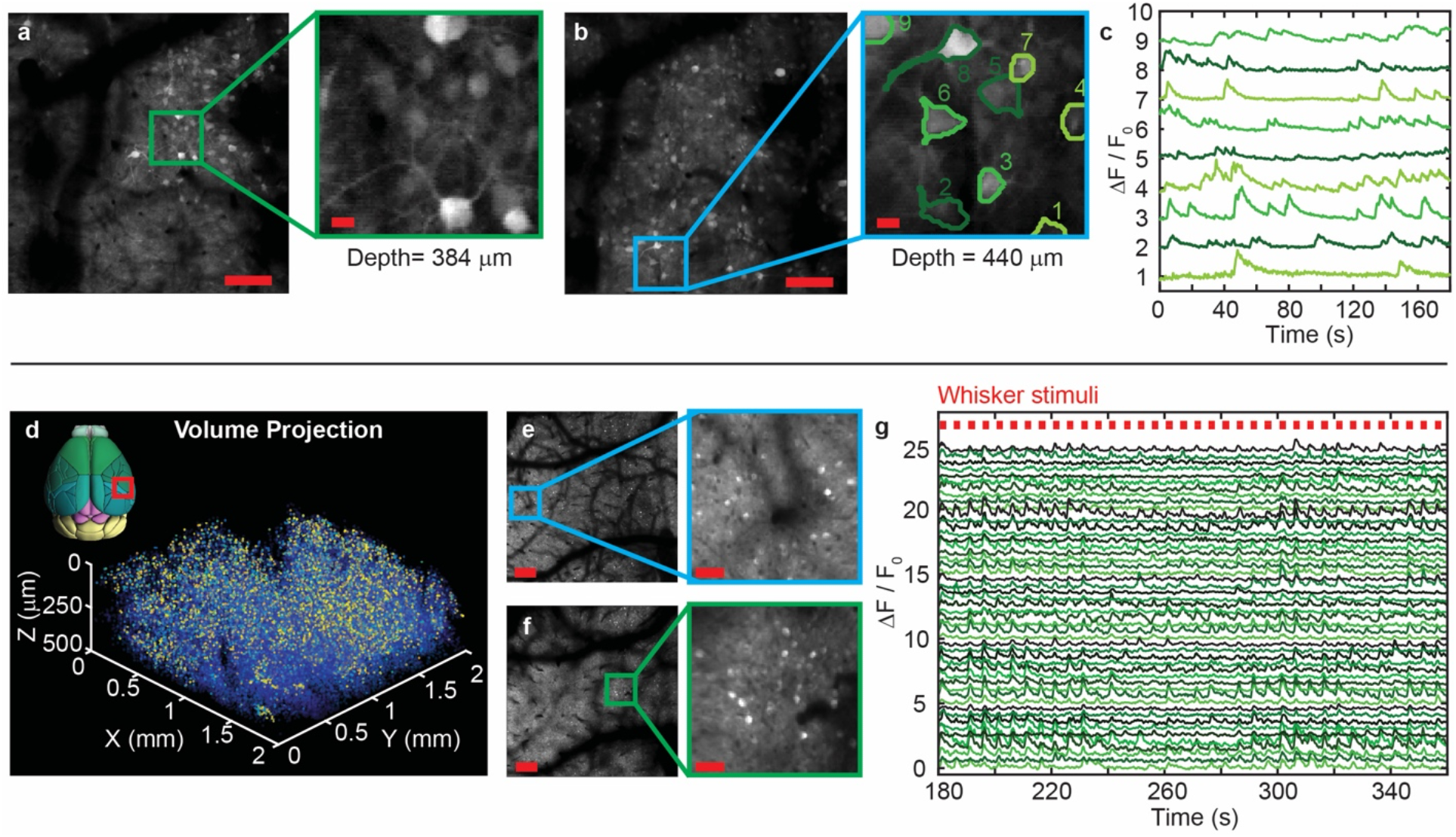
Multi-scale functional imaging with light beads microscopy. **a–c,** High resolution volumetric (~600 × 600 × 500 μm^3^) imaging of neuroactivity at 9.6 Hz in jGCaMP7f-expressing mice. Representative mean projection images of neurons at planes **a** 440 μm and **b** 384 μm deep, respectively taken from the above volume during a 3-minute recording. Scale bars: 50 μm. Zoomed-in regions inset, scale bars: 10 μm. See also Movie S6. **c,** Representative timeseries of the 9 neurons outlined in the zoomed-in region of the plane in **b.** Offset: 1.0 *ΔF/F_0_*. **d–g,** Recording of 70,275 neurons within a volume of ~2 × 2 × 0.5 mm^3^ at 6.7 Hz and 2.8 μm lateral voxel sampling. **d,** 3D rendering of extracted neuron spatial coordinates and maximum-projected activity for a 9-minute recording. Transverse brain image reproduced from Ref. 52. See also Movie S7. **e,f** Mean projection images at 144 and 482 μm depths, respectively. Scale bars: 250 μm. Zoomed-in regions inset, scale bars: 50 μm. See also Movie S8. **g,** Representative time-series of 50 whisker-tuned neurons. Occurrences of the stimulus denoted by red marks. Offset: 0.5 *ΔF/F_0_*.

Finally, by employing our optimized ~5 μm lateral voxel sampling we can image a volume of ~5.4 × 6.0 × 0.5 mm^3^ encompassing both hemispheres of the mouse neocortex down to a depth of ~600 μm in tissue (Fig. 5, Movies S9 and S10, 5.4 × 6 × 0.5 mm^3^ FOV). In this modality, we demonstrate Ca^2+^imaging of populations of 0.8–1.1 million neurons at single neuron resolution and 2.2 Hz volume rate (3 recordings, N = 3 mice, Fig. S6b). Fig. 5 shows a representative recording of 807,748 neurons at which single calcium transients can be clearly detected (Fig. 5d, Movie S11). The optical access, large degree of multiplexing, and optimized scanning approach employed by LBM opens the door to scaling 2pM from single-brain-regions to cortex-wide recording, allowing for investigation of bi-hemispheric cognitive processing, as well as capturing the single trial dynamics of populations of neurons in the mammalian brain more than ~2 orders of magnitude larger than any other technique.^10,23,25,29^

**Fig. 5:**
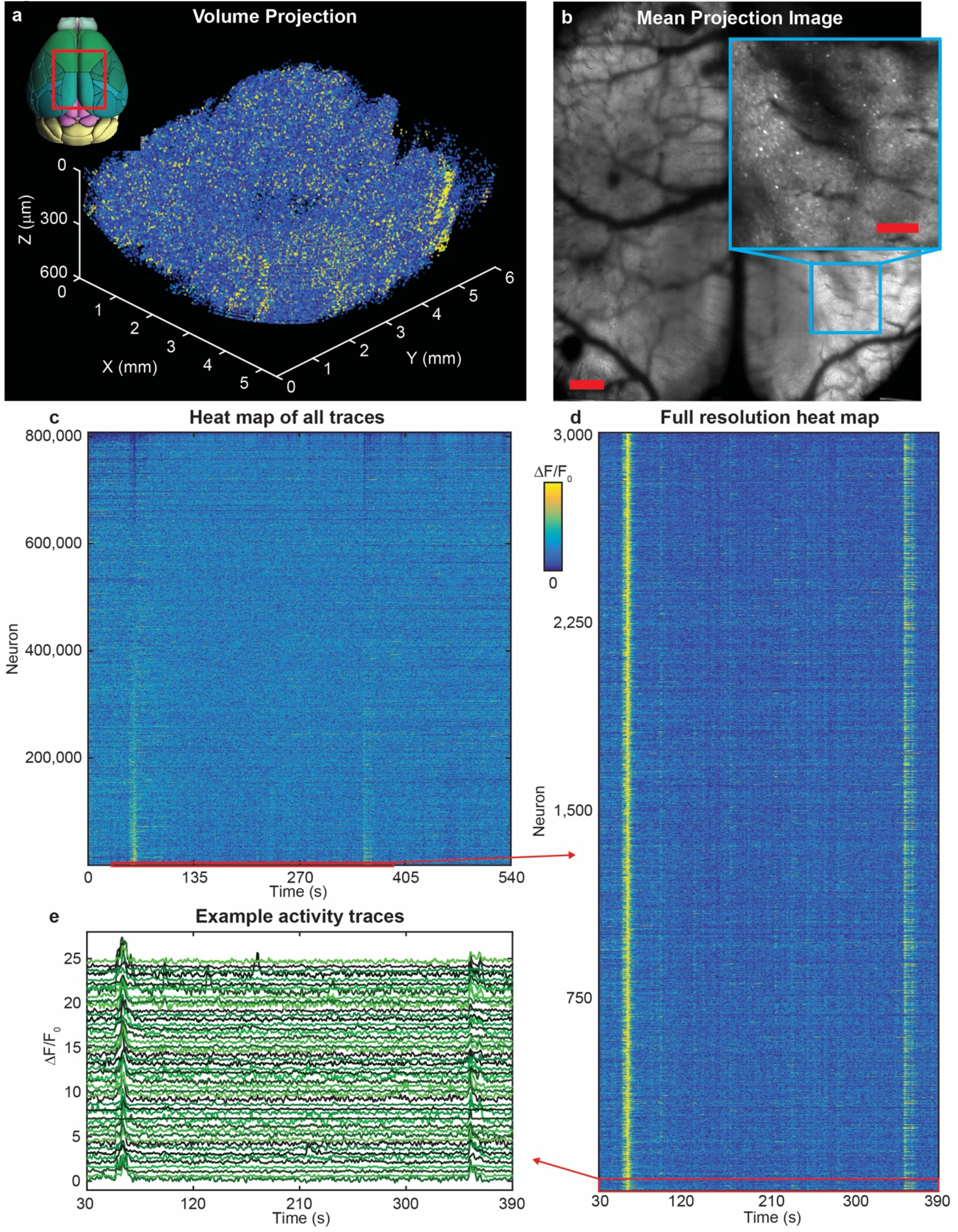
Volumetric recording of 807,748 neurons within ~5.4 × 6 × 0.5 mm^3^ volume at 2.2 Hz in a GCaMP6s-expressing mouse. **a,** 3D rendering of extracted neuron spatial coordinates and maximum projected activity for a 9-minute recording. Transverse brain image reproduced from Ref. 52. See also Movie S9. **b,** Mean projection image from the recording in a at 600 μm depth. Scale bar: 500 μm. Inset: zoomed in region of **b**, scale bar: 200 μm. See also Movie S10. **c,** Heat map of all 807,748 extracted neurons normalized to maximum *ΔF/F_0_*. See also Movie S11. Subset of 10 example traces. Offset: 1.0 *ΔF/F_0_*. **d,** Subset of 3,000 traces from **c** shown at full resolution. **e,** Subset of 50 traces from **d**. Offset: 0.5 *ΔF/F_0_*.

## Discussion

Mesoscopic 2pM platforms are a necessary tool for increasing the optical access of functional calcium imaging to multi-regional recording from the mammalian brain. Here we have argued that operating at the optimal condition for spatiotemporal sampling is essential for pushing the combined tradeoff between speed, volume size and resolution to the limits that allows designing an imaging system capable of capturing calcium transients from single neurons throughout the mouse cortex (see Supplementary Note 1). Maximizing efficiency of excitation and volumetric recording speed requires one-pulse-per-voxel, SNR-maximized excitation, continuous voxel acquisition at fluorescence-lifetime-limited rates, and spatial sampling at the minimum density required to resolve cells or other features of interest. This in turn frees up resources such as the brain-heating-limited power budget and excess SNR that can be used to further increase volume size, speed or resolution.

LBM represents the first realization of the optimal condition described above for spatiotemporal acquisition. This is enabled by the MAxiMuM module of our LBM which allows for scaling of the multiplicity of temporal multiplexing in the axial direction beyond the previously shown few-beam regime, making full use of the pulse-to-pulse time interval of our laser source in order to achieve the maximum voxel rate possible within the fluorescence lifetime limit of GCaMP. Crucially, the cavity-based design of MAxiMuM allows for an axial multiplexing scheme in which the transmission of the cavity can be tuned to flexibly accommodate the need for increasing power to maintain constant SNR with each subsequent beam focused deeper in the tissue. This is in contrast to previous axial multiplexing methods where inherent design constraints require wide spacing of the axial planes (>100 μm) to maintain SNR and thus limits multiplicity to only a handful of planes possible within the penetration depth of 2pM.^34^ Furthermore, in comparison to MAxiMuM, other temporal multiplexing techniques have been limited to non-mesoscopic^19,23,25,30,33^ or non-volumetric^10,19,23,30-33^ imaging due to a combination of inefficiencies including the unfavorable scaling of multiplicity with the system’s complexity, incompatibility with axial multiplexing, or incompatibility with one-pulse-per-voxel excitation. In contrast, LBM, in its myriad configurations, operates at fluorescence-lifetime-limited voxel acquisition rates, in one-pulse-per-pixel mode, and with a high degree of multiplicity which together enable optimal voxel sampling for volumetric recording at the highest possible voxel acquisition rate. Thereby, compared to existing techniques, LBM allows for up to an order of magnitude effective increase in the total number of accessed voxels, and a 1-2 order of magnitude increase in recording volume, scaling the total number of recorded neurons by up to 3 orders of magnitude, while maintaining calcium-imaging-compatible frame rates (Table 1).

**Table 1:** Summary of LBM recording modalities in comparison to existing multiplexed imaging techniques in the literature.

| Specifications: | Imaging modality: |  |  |  |  |  |  |  |
| --- | --- | --- | --- | --- | --- | --- | --- | --- |
|  | Single hemisphere LBM | High-resolution LBM | High-speed LBM | Bi-hemispheric LBM | HyMS | FACED | Reverberation Microscopy | 16 beam microscopy |
| 1 pulse-per-voxel [yes/no] | Yes | Yes | Yes | Yes | Yes | No | Yes | No |
| Fluorescence lifetime limited [yes/no] | Yes | Yes | Yes | Yes | No | No | No | No |
| Axial or lateral multiplexing | Axial | Axial | Axial | Axial | Axial or lateral | Lateral | Axial | Lateral |
| Multiplicity | 30 | 30 | 30 | 30 | 4 | 80 | 4-6 | 16 |
| Volumetric [yes/no] | Yes | Yes | Yes | Yes | Yes | No | No | No |
| FOV ($x \times y \times z$) [mm<sup>3</sup>] | $3 \times 5 \times 0.5$ | $0.6 \times 0.6 \times 0.5$ | $0.9 \times 0.9 \times 0.5$ | $5.4 \times 6 \times 0.5$ | $1.0 \times 1.0 \times 0.6$ | Single plane: $0.05 \times 0.25$ | 4 axial planes: $0.6 \times 0.6$ | Single plane: $2.0 \times 2.0$ |
| Lateral sampling ($x \times y$) [μm] | $4.2 \times 5$ | $1 \times 1$ | $3.1 \times 3.1$ | $4.2 \times 5$ | $5 \times 5$ | $0.23 \times 0.6$ | Unspecified | $1.25 \times 1.25$ |
| Total number of voxels accessed | $25.9 \times 10^6$ | $10.4 \times 10^6$ | $2.6 \times 10^6$ | $46.7 \times 10^6$ | $2.7 \times 10^6$ | $72 \times 10^3$ | Unspecified | $2.6 \times 10^6$ |
| Voxel acquisition rate [MHz] | 141 | 141 | 141 | 141 | 18.8 | 80 | 80-120 | 47 |
| Volume rate [Hz] | 4.7 | 9.6 | 38 | 2.2 | 5.1 | 1,000 | 30 | 18.5 |
| Number of neurons recorded | ~200,000 | ~4,000 | - | ~1,000,000 | ~12,000 | < 100, estimated from Fig. 2b | < 100, estimated from Fig. 3a | ~2,000 |
| Brain regions imaged simultaneously | SSp, VISp, PTLp, RSP | PTLp | - | SSp, VISp, PTLp, RSP, bilaterally | AUDp | VISp | MO or OLF | VISp |
| Reference | Fig. 2 | Fig. 4a-4c | - | Fig. 5 | Ref. 25 | Ref. 33 | Ref. 34 | Ref. 10 |

Enabled by these capabilities of LBM, our approach has uniquely allowed finding, amongst other observations, evidence for mixed selectivity^42^ in large populations of neurons distributed across many regions of the brain, as well as trial-to-trial variability of both stimulus- and behavior-tuned neurons. Additionally, we found evidence for covariance of activity amongst subsets of the stimulus-tuned neuronal population across the brain at the single-trial level which have been suggested to represent encoding of information about internal states and uncontrolled aspects of stimulus and behavior.^11,40^ These observations highlight the need for both high-speed and large-scale neuronal recording capability in order to identify and capture long-range functional cortical circuits and the variability of their response at the trial-to-trial level and single neuron resolution. Moreover, as suggested by our observations on the correlation distance for the simple sensory and behavioral paradigms used in this work (2 – 4 mm), and bolstered by other findings in recent literature,^9,16,40^ mesoscopic-scale volumetric imaging of population sizes on the order of 10^5^ – 10^6^ neurons is necessary for revealing the full neuronal population code in individual cortical regions,^9,16,40,43^ as well as identifying the structure and dynamics^11,13,44-46^ of inter-regional brain activity critical for learning,^12,47^ memory,^14,48^ and other cognitive functions. As such, the unprecedented size of neuronal population recording enabled by our technique opens up a new range of opportunities to understand how the neurocomputations underlying multi-regional encoding and processing of sensory and behavioral information emerge from the dynamic interaction of brain-wide networks of neurons at the single-neuron level in the mammalian brain.

## Methods

### Laser source

Our custom laser system was comprised of an ultrafast ytterbium-doped fiber master oscillator and chirped-pulse amplifier (Active Fiber Systems, 60 W average power, 4.7 MHz, 300 fs pulse duration, ~10 μJ pulse energy, λ = 1030 nm), followed by an optical parametric chirped-pulse amplifier (OPCPA, White Dwarf dual, Class5 Photonics). The OPCPA operated at a wavelength of 960 nm with ~90 fs duration pulses up to ~0.8 μJ in energy at a repetition rate of 4.7 MHz. We employed an electro-optic modulator (Conoptics, 350-160-BK) to dynamically adjust laser power in the sample and blank the beam while the resonant scanner was reversing direction. We precompensated pulse-broadening using two pairs of chirped mirrors (Class5 Photonics) with −500 fs^2^ per reflection which imparted a total of −24,000 fs^2^ of anomalous group delay dispersion to counteract the material dispersion of the multiplexing module, the mesoscope, and the other components in the system.

### Spatiotemporal multiplexing module

To facilitate spatiotemporal multiplexing, we constructed a cavity comprised of concave mirrors configured in an 8-f, non-inverting, re-imaging scheme (Fig. S1a). The input beam was focused by L_1_, just above the aperture of the partially-reflective mirror (PRM), M_1_, and in the front focal plane of M_2_. Mirrors M_2_–M_5_ were concave mirrors (f = 500 mm, 2” diameter) with custom low-dispersion dielectric coatings (Layertec, Inc) which re-imaged the initial spot onto the turning mirror M_6_. M_6_ provided a slight vertical tilt to the beam such that it intersected the PRM M_1_. M_1_ was a low dispersion ultrafast beam splitter (Thorlabs, UFBS9010) with a nominal transmission of ~10% at 45° incidence. By adjusting the position of M_6_, we were able to change the angle of incidence at the PRM and tune the transmission to the desired value of ~8%. The majority of the light incident on M_1_ underwent the next round-trip through the cavity, and the rest of the light was transmitted. Each round trip through the cavity provided a temporal delay *τ* = 13.8 ns, as well as an offset in the focal plane of the beam, dictated by the distance between M_6_ and M_1_ (~145 mm). The vertical angle of M_6_, necessary to ensure the beam intersected the aperture of M_1_, caused a small lateral offset between subsequent round trips. This offset was minimized during alignment to reduce the offset between axial planes in the sample. Round trips in the primary cavity generated the first 15 multiplexed beams, and a subsequent single-pass cavity (Fig. S1b) increased the multiplicity to 30.

After the primary cavity (Fig. S1a), the light was re-collimated by L_2_. L_3_ and L_4_ formed a unitary magnification telescope that ensured that the lowest power beams were directed to the shallowest depths in the sample. The distances between M_6_ and L_2_, L_2_ and L_3_, and L_3_ and L_4_ were iteratively optimized in order to position the last beam exiting Cavity A in the nominal focal plane of the objective, while maintaining as uniform as possible magnification for each beam. The beams were transmitted through a half-wave plate (HWP) and onto a polarizing beam splitter (PBS). The reflected portion of the beam underwent a single round-trip through another custom-mirror-based 8-f re-imaging cavity (f = 250 mm, 2” diameter, Layertec, Inc), before recombination with the transmitted portion of the beam (Fig. S1b). The beams coupled to the secondary cavity were delayed an additional 6.7 ns, interleaving them in time with the beams transmitted by the PBS (Fig. S1c). The focal planes of these delayed beams could be globally shifted by adjusting the position of M9 and M11, and formed two sub-volumes that were spatially contiguous, such that all 30 beams provided continuous sampling along the optical axis (Fig. S1d). Manipulation of the HWP could be used to adjust the relative optical power of the sub-volumes in order to preserve matching to the scattering properties of the tissue. In total, 30 spatiotemporally multiplexed beams exited the secondary cavity, and the axial separation between imaging planes is ~16 μm leading to a total axial sampling range of 465 μm.

### Integration with mesoscope

The output of the multiplexing module was interfaced with a commercial mesoscope (Thorlabs, Multiphoton Mesoscope).^18^ The mesoscope layout and accompanying electronics are shown in Fig. S2. The configuration of the microscope followed normal operation conditions with the exception of some minor modifications. The remote focusing unit of the system, consisting of a PBS and objective mounting apparatus, was removed and replaced by a turning mirror to route beams directly to the first telescopic relay. This modification was necessary because light exiting MAxiMuM was split between two orthogonal polarization states and thus incompatible with the PBS in the remote focusing module. Furthermore, the axial range of MAxiMuM (~500 μm) makes remote focusing redundant for our intended axial imaging range and thus an unnecessary drain on the power and dispersion compensation budgets.

Additionally, the electrical amplifier following the photo-multiplier tube (PMT) was removed, as the temporal response of the standard model amplifier used with the mesoscope was insufficient for multiplexed data. Signals in the one-pulse-per-voxel operation regime are much higher than those for conventional pulse-averaging detection modalities; for example, signals in our system were 17× higher than that of a typical 80 MHz pulse train with equivalent average power. Accordingly, and consistent with our previous demonstrations,^25^ we found that additional amplification was not necessary.

### Data acquisition

Data was acquired using the commercial mesoscope-compatible version of the ScanImage software platform (Vidrio, Inc.) with some additional customizations, as well as upgraded digitization hardware (Fig. S2a). We used an evaluation board (Analog Devices, AD9516-0) to multiply a trigger signal from the OPCPA laser to 1614 MHz, which in turn was fed to the upgraded digitizer (National Instruments, NI 5772) and field programmable gate array (FPGA, National Instruments, PXIe-7975R) to serve as a sample clock. This clock signal was used within the customized version of ScanImage to synchronize the line trigger to the pulse repetition rate of the laser, thus ensuring a single laser pulse constituted one voxel of the recording.

Additionally, the ScanImage customization allowed the user to define channels by integrating temporal windows of the raw PMT signal (Hamamatsu H11706-40) with respect to a trigger from the laser. The window for each channel was set to integrate the fluorescence signal associated with each beam from the MAxiMuM system such that the channels constitute the de-multiplexed axial planes of the volumetric recording (channel plots in Fig. S2b). The microscope recorded frames for each channel separately, in the same fashion as a two-color compatible microscope records separate channels from each PMT. Data streamed to disk consisted of 30 consecutive frames representing each channel, and thus each axial plane, repeated in sequence for each time point in the measurement.

### Data processing

Figure S4 shows a schematic of the data processing pipeline. Data recorded by the microscope was reassembled from constituent ROIs into 30 stacks of frames (*x,y,t*) corresponding to each plane of the volume which were each processed separately. Motion correction of each plane was facilitated using the non-rigid version of the NoRMCorre algorithm^49^ and neuronal footprints and time-series were extracted with using the planar, patched version of the CaImAn software package.^50,51^ Due to the reduced spatial resolution of the data, the elliptical search method was found to most accurately extract neuronal signals from soma. The algorithm was initialized with a number of components dictated by the physiological expectation from the given volumetric field of view, assuming a standard density of 9.2×10^4^ neurons per cubic millimeter.^52,53^ The spatial correlation threshold was held at the default value of 0.4, and the minimum signal-to-noise parameter was set to 1.4. In practice we found that this value was consistent with only keeping transients with statistically significant transient activity (see statistics in the following section). Finally, neuronal footprints were screened using the ‘mn’ and ‘mx’ options in CaImAn such that components larger than the area expected for a 20 μm diameter neuron, or smaller than that of a 10 μm diameter neuron, in the equivalent pixel space, were eliminated.

The detected neurons from each plane in the volume were subsequently collated. The lateral positions of neuronal footprints were corrected for plane-to-plane offset using calibration values determined by recordings of pollen grains (Fig. S3d). For cases where components in adjacent planes were temporally correlated above the default CaImAn threshold (0.8) and also had any spatially overlapping voxels, the time-series and footprints were merged into a single component. First order moments in the *x, y*, and *z* directions were used to determine the centroids of each neuronal component. The field curvature imposed by the microscope was corrected using a parabolic profile with a – 158 μm offset at the periphery of the full FOV, following the characterization in Ref. 18.

### Data analysis

Correlations between neuronal activity and stimuli were analyzed by correlating the time series of each neuron with the corresponding stimulus vector, generated by convolving a time-series composed of the onset of each stimulus or behavior with the expected kernel of the calcium indicator (see the final panel of Fig. S4). This kernel had an exponential rise time of 200 ms and a decay of 550 ms in agreement with the literature values for GCaMP6s.^5^ All correlations considered between stimulus vectors and neuronal time-series were Pearson type and used the raw time-series data rather than the deconvolved traces from CaImAn. The lag between the neuronal time-series and each stimulus vector was defined as the time for which the cross-correlation between each trace and vector was maximized. For determining stimulus-tuned populations (Figs. 3b–3d and S7a–7c), the median value of the distribution of lags was applied as an offset to each time-series prior to determining correlation. For the temporal analysis in Figs. 3r and 3s, the relative lag values for each individual behavior-tuned neuron are shown with respect to the median lag value.

Null hypothesis testing was conducted by creating a time-series with a number of randomly shuffled “stimuli” equal to the number presented during a typical recording. For uninstructed behaviors, shuffling was achieved by circulating each trace in the data set by a random value to remove temporal structure. The threshold for significant correlation with visual stimuli, whisker stimuli, or uninstructed animal behaviors was determined by fitting the shuffled correlations, *r*, to a normal distribution given by 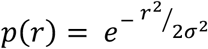. Correlations with stimuli for which *r* > 3*σ* were considered highly correlated, while correlations below *σ* were deemed insignificant.

Hierarchical clustering was performed using Ward’s method with the Euclidean distance metric via the MATLAB function ‘linkage.’ For the mixed representation analysis in Figs. 3j–3o, the activity of each trial was defined as the integration of the time-series of each neuron in a 5 second window following the presentation of the stimulus. Significance of the change in a neuron’s activity was determined with a t-test comparing the activity of all trials with the stimulus presented alone to those where the stimulus was presented coincidently with another stimulus. Neurons with *p < 0.05* were considered to have significant change in activity.

### Animal statistics

A total of *N* = 6 animals transgenically-expressing GCaMP6s, and *N* = 3 animals expressing jGCaMP7f^41^ through viral transfection were imaged during experiments (VGlut-IRES-Cre driver, Ai162 reporter).^5,37^ Figures S6c–6e show distributions of typical transient activity in GCaMP6s-expressing mice, including peak activity, baseline noise levels, and typical transient decay times. The characterizations are consistent with expectations for two photon imaging. Figure S6f shows the distribution of maximum Z-scores for the neuronal data set shown in Fig. 2. Z-scores were calculated by applying a 3-point moving average to each neuron’s time-series, finding the maximum value and normalizing by the baseline noise. The moving average in this instance ensures that we are measuring the robustness of all consecutive data points within the kernel of the indicator (200 ms rise time and 550 ms decay time for GCaMP6s sampled at a 4.7 Hz volume rate implies 3 samples within each transient) relative to noise, and not inflating the significance of isolated fluctuations in the data. At an SNR threshold of 1.4, the cutoff of the distribution is such that neurons in the data set have activity exceeding at least three standard deviations of the baseline (Fig. S6f), indicating low likelihood of false-positive ROIs being classified as neurons. Figure S6g shows the distribution of nearest neighbor separations between neurons in the data set. The majority of pair-wise distances occur between 10 – 20 μm, in agreement with expectations for cortical neuronal density.^5,37^

### Apparatus for stimulus delivery and behavioral tracking

Visual and somatosensory stimuli were controlled via a synchronization script running in parallel to ScanImage implemented on a microcontroller (Arduino Uno). A portion of the voltage used to open the laser shutter was read in by the microcontroller, triggering the beginning of a recording epoch and synchronizing the microcontroller clock to the ScanImage frame clock. For whisker stimulation, a motor shield and servo motor were employed to move a brush forward and backward over the animals’ whiskers at time intervals indicated by the stimulation protocol. The brush size and its proximity were chosen to stimulate all whiskers simultaneously (as opposed to stimulation of specific whiskers), and stimulation was applied contralaterally to the hemisphere being recorded by the microscope.

For visual stimuli, the microcontroller sent a 5 V TTL trigger signal to the control computer. A parallel MATLAB program read in these trigger signals and generated a series of images on a secondary external monitor (Waveshare 11.6” LCD) placed ~20 cm from the animal’s eyes. For each trigger signal, a 500 ms duration movie was displayed on the monitor consisting of a binary drifting grating pattern at the full dynamic range of the screen. The position of the screen was chosen to cover 72° of the animal’s field of view horizontally and 43° vertically. The grating period was 0.07 cycles per degree, and the rate of drift was 1 cycle/second. The orientation of the grating followed a pattern of 0° (horizontal), 45°, 90° (vertical), and 135° and was repeated in this pattern for all stimuli during the recordings.

All rodents were head-fixed on a home-built treadmill with a rotation encoder affixed to the rear axle (Broadcom, HEDS-5540-A02) to measure the relative position of the tread during the recordings. Treadmill position, the microcontroller clock value, and the onset of either a visual or whisker stimulus were streamed to the control computer via a serial port connection and logged with a separate data logging script. The data logging script also triggered a camera (Logitech 860-000451) in order to capture additional animal behavior during recordings. Motion tracking of the rodent’s left and right forelimbs and right hindlimb was facilitated using DeepLabCut.^54,55^ An example recording of the animal with motion tracking super-imposed over top is shown in Movie S1. Stimulation was presented at 5 second intervals such that the calcium signal from correlated transients had sufficiently decayed before the onset of the next stimulus.

### Data visualization

All time-series data is displayed with a moving average corresponding to a 1 s time interval along the temporal axis to improve transient visibility. Calcium trace heatmaps are individually normalized to improve visualization. For 3D visualization, equally sized spheres in the data set were rendered using the ‘scatter3’ function in MATLAB. For Movies S2, S7, and S9, the top ~57,000, ~33,000, and ~150,000 most-active neurons, respectively, are visualized and their time-series are individually normalized, with the opacity of the representative sphere increasing with transient activity. For the volume projection images in Figs. 2a, 4d, and 5a in the manuscript, the top ~207,000, ~70,000, and ~202,000 most-active neurons are rendered, respectively, and the color of each sphere represents the maximum projection of the corresponding neuron’s time-series with the color bar and opacity of each representative sphere adjusted for maximum visibility of the most active neurons.

### Imaging power and immunohistochemical validation

We used 150 – 200 mW of power to image FOVs of 0.6×0.6×0.5 mm with sufficient signal-to-noise ratio in jGCaMP7f-expressing mice. In transgenic mice, power was restricted to < 250 mW in small FOVs (0.6×0.6×0.5 mm, 2×2×0.5 mm) to remain within previously established thresholds for brain safety. For large FOV recordings (3×5×0.5 mm, 5.4×6×0.5 mm), power ranged from 200-450 mW.

To further validate any possible neuropathological responses associated with these intensities and absolute power levels when delivered within the large volumetric FOVs and cranial windows, we employed immunihistochemical labeling. Brain sections were immunostained for astrocyte activation marker GFAP following imaging under various laser intensities. All experiments were conducted at least 2 weeks after cranial window surgery. Awake head-fixed mice (wild-type) were subject to various laser powers and scanning FOVs (see Fig. S8) for 9 minutes continuous at a depth of ~600 um below the surface of the brain. For verification, we included a negative control condition corresponding to animals that had undergone cranial window implantation but had not been exposed to laser power in the region of the brain considered. As a positive control, we imaged with 360 mW of power in a FOV of 0.4 mm, exceeding previously established limits for brain safety.^24^ To make full use of the 8 mm cranial window and each animal, both hemispheres of each mouse were used for separate experiments, with negative control and low exposure conditions contralateral to high exposure and positive control conditions. 16 h after scanning, mice were deeply anaesthetized with isoflurane (4% flow rate of 0.5–0.7 l/min) and transcardially perfused with cold phosphate-buffered saline (PBS) followed by 4% PFA (VWR International Inc, 15710). Brains were extracted and placed in PFA for 24 h and then transferred to 30% sucrose/PBS solution at 4 °C. Coronal sections (30 μm thickness) were collected from within and around the scanning FOV site using a cryostat (Leica Biosystems). Brain sections were permeabilized using 0.2% Triton-X100/PBS (PBST) for a 1 h incubation period, followed by a blocking solution of 5% normal goat serum (NGS) in PBST for 1 h. Sections were then incubated in primary mouse GFAP antibody (Protein Tech, 60190-1-Ig) (1:800) in PBST + 2% NGS for 24 h at 4 °C. Sections were then washed three times with PBS for 20 minutes per wash, followed by an incubation period in Alexa 594-conjuagted goat anti-mouse antibody (Abcam, ab150116) (1:1000) for 2 h at room temperature. Sections were washed again three times with PBS for 20 minutes per wash with Hoechst 33342 (Invitrogen, H3570) (1:2000) being added during the last wash, before being mounted on slides and coverslipped using anti-fade mounting medium (Invitrogen, P10144).

Brain sections were imaged at 20× magnification using a resonant-scanning confocal microscope (Caliber I.D, RS-G4). Images were analyzed using FIJI. Relative fluorescent intensity was quantified by measuring the mean fluorescent intensity in a 1×1 mm axial area centered within the imaging FOV and dividing this measurement by the mean intensity of equivalently sized areas within the control hemispheres.

### Brain heating simulations

We used a finite difference model^56^ to simulate laser-induced heating, thermal conductivity, and homeostatic cooling through blood perfusion. Additionally, we employed modifications^57^ account for a scanned focal plane and heat conduction through the cranial window and immersion water. All simulations used an 8 mm cranial window, with the exception of those in Fig. S8j where the window size varies from 3 to 8 mm. The boundary conditions of the model were adjusted to assume a constant temperature of 25 °C at a distance 1.5 mm above the surface of the cranial window. We used a voxel size of 0.01 mm for light diffusion and 0.03 mm for heat diffusion, a time step of 0.16 ms and an optical wavelength of 960 nm. Material constants for glass and water were obtained from published tables.

## Acknowledgments

We thank P. Strogies and J. M. Petrillo at the Precision Instrumentation Technology (PIT) of the Rockefeller University for manufacturing mechanical components; S. Weisenburger for helpful discussions related to microscope development and synchronization; and T. Nöbauer for discussions regarding data management and processing.

## Contributions

J. D. contributed to the project conceptualization, designed and built the imaging and data acquisition system, performed experiments, programmed experimental control and analysis software, analyzed data, and wrote the manuscript. J. M. performed data processing and modeling and contributed to writing the manuscript. F. T. contributed to microscope construction and characterization and aided in developing the experimental control design. H. K., F. M. T., and B. C. performed virus injections and cranial window surgeries. A.V. conceived and led the project, designed the imaging system, the data acquisition approach, and all *in vivo* mouse experiments, and wrote the manuscript.

## References

1. W. Denk, J. H. Strickler, W. W. Webb, Two-photon laser scanning fluorescence microscopy. Science 248, 73–76 (1990).

2. P. T. C. So, C. Y. Dong, B. R. Masters, K. M. Berland, Two Photon Excitation Fluorescence Microscopy. Annu. Rev. Biomed. Eng. 2, 399–429 (2000).

3. F. Helmchen, W. Denk, Deep tissue two-photon microscopy. Nat. Methods 2, 932–940 (2005).

4. A. Miyawaki, J. Llopis, R. Heim, J. M. McCaffery, J. A. Adams, M. Ikura, R. Y. Tsien, Fluorescent indicators for Ca2+ based on green fluorescent proteins and calmodulin. Nature 388, 882–887 (1997).

5. J. Nakai, M. Ohkura, K. Imoto, A high signal-to-noise Ca(2+) probe composed of a single green fluorescent protein. Nat. Biotechnol. 19, 137–141 (2001).

6. T. Chen, T. Wardill, Y. Sun, S. R. Pulver, S. L. Renninger, A. Baohan, E. R. Schreiter, R. A. Kerr, M. B. Orger, V. Jayaraman, L. L. Looger, K. Svoboda, D. S. Kim, Ultrasensitive fluorescent proteins for imaging neuronal activity. Nature 499, 295–300 (2013).

7. J. A. Harris, S. Mihalas, K. E. Hirokawa, J. D. Whitesell, H. Choi, A. Bernard, P. Bohn, S. Caldejon, L. Casal, A. Cho, A. Feiner, D. Feng, N. Gaudreault, C. R. Gerfen, N. Graddis, P. A. Groblewski, A. M. Henry, A. Ho, R. Howard, J. E. Knox, L. Kuan, X. Kuang, J. Lecoq, P. Lesnar, Y. Li, J. Luviano, S. McConoughey, M. T. Mortrud, M. Naeemi, L. Ng, S. W. Oh, B. Ouellette, E. Shen, S. A. Sorensen, W. Wakeman, Q. Wang, Y. Wang, A. Williford, J. W. Phillips, A. R. Jones, C. Koch, H. Zeng, Hierarchical organization of cortical and thalamic connectivity. Nature 575, 195–202 (2019).

8. K. V. Kuchibhotla, J. V. Gill, G. W. Lindsay, E. S. Papadoyannis, R. E. Field, T. A. Hindmarsh Sten, K. D. Miller, R. C. Froemke, Parallel processing by cortical inhibition enables context-dependent behavior. Nat. Neurosci. 20, 62–71 (2017).

9. C. Stringer, M. Pachitariu, N. Steinmetz, M. Carandini, K. D. Harris, High-dimensional geometry of population responses in visual cortex. Nature 571, 361–365 (2019).

10. O. I. Rumyantsev, J. A. Lecoq, O. Hernandez, Y. Zhang, J. Savall, R. Chrapkievicz, J. Li, H. Zeng, S. Gaguli, M. J. Schnitzer, Fundamental bounds on the fidelity of sensory cortical coding. Nature 580, 100–105 (2020). NC

11. S. Musall, M. T. Kaufman, A. L. Juavinett, S. Gluf, A. K. Churchland, Single-trial neural dynamics are dominated by richly varied movements. Nat. Neurosci. 22, 1677–1686 (2019). NC

12. H. Makino, C. Ren, H. Liu, A. N. Kim, N. Kondapeneni, X. Liu, D. Kuzum, T. Komiyama, Transformation of Cortex-wide Emergent Properties during Motor Learning. Neuron 94, 880–890 (2017).

13. T. Mao, D. Kosefoglu, B. M. Hooks, D. Huber, L. Petreanu, K. Svoboda, Long-Range Neuronal Circuits Underlying the Interaction between Sensory and Motor Cortex. Neuron 71, 111–123 (2011).

14. L. Pinto, K. Rajan, B. DePasquale, S. Y. Thiberge, D. W. Tank, C. D. Brody, Task-Dependent Changes in the Large-Scale Dynamics and Necessity of Cortical Regions. Neuron 104, 810–8214 (2019).

15. Q. Lin, J. Manley, M. Helmreich, F. Schlumm, J. M. Li, D. N. Robson, F. Engert, A. Schier, T. Nöbauer, A. Vaziri, Cerebellar Neurodynamics Predict Decision Timing and Outcome on the Single-Trial Level. Cell 180, 1–16 (2020).

16. R. Bartolo, R. C. Saunders, A. R. Mitz, B. B. Averbeck, Information-Limiting Correlations in Large Neural Populations. J. Neurosci. 40, 1668–1678 (2020).

17. P. S. Tsai, Celine Mateo, Jeffrey J. Field, Chris B. Schaffer, Matthew E. Anderson, David Kleinfeld, Ultra-large field-of-view two-photon microscopy. Opt. Express 23, 13833–13847 (2015).

18. N. J. Sofroniew, D. Flickinger, J. King, K. Svoboda, A large field of view two-photon mesoscope with subcellular resolution for in vivo imaging. eLife 5, 14472 (2016).

19. J. N. Stirman, I. T. Smith, M. W. Kudenov, S. L. Smith, Wide field-of-view, multi-region, two-photon imaging of neuronal activity in the mammalian brain. Nat. Biotechnol. 34, 857–862 (2016).

20. N. Ji, J. Freeman, S. L. Smith, Technologies for imaging neural activity in large volumes. Nat. Neurosci. 19, 1154–1164 (2016).

21. S. Weisenburger, A. Vaziri, A Guide to Emerging Technologies for Large-Scale and Whole-Brain Optical Imaging of Neuronal Activity. Annu. Rev. Neurosci. 41, 431–452 (2018).

22. W. Yang, R. Yuste, In vivo imaging of neural activity. Nat. Methods 14, 349–359 (2017).

23. C. Yu, J. N. Stirman, Y. Y. Riichiro Hira, S. L. Smith. https://doi.org/10.1101/2020.09.20.305508(2020).

24. R. Prevedel, A. J. Verhoef, A. J. Pernía-Andrade, S. Weisenburger, B. S. Huang, T. Nöbauer, A. Fernández, J. E. Delcour, P. Golshani, A. Baltuska, A. Vaziri, Fast volumetric calcium imaging across multiple cortical layers using sculpted light. Nat. Methods 13, 1021–1028 (2016).

25. S. Weisenburger, F. Tejera, J. Demas, B. Chen, J. Manley, F. T. Sparks, F. M. Traub, T. Daigle, H. Zeng, A. Losonczy, A. Vaziri, Volumetric Ca2+ Imaging in the Mouse Brain using Hybrid Multiplexed Sculpted Light (HyMS) Microscopy. Cell 177, 1–17 (2019).

26. A. Kazemipour, O. Novak, D. Flickinger, et al., Kilohertz frame-rate two-photon tomography. Nat Methods 16, 778–786 (2019).

27. E. J. Botcherby, R. Juškaitis, T. Wilson, Scanning two photon fluorescence microscopy with extended depth of field. Opt. Commun. 268, 253–260 (2006).

28. A. Song, A. Charles, S. Koay, J. L. Gauthier, S. Y. Thiberge, J. W. Pillow, D. W. Tank, Volumetric two-photon imaging of neurons using stereoscopy (vTwINS). Nat. Methods 14, 420–426 (2017).

29. R. Lu, Y. Liang, G. Meng, P. Zhou, K. Svoboda, L. Paninski, N. Ji, Rapid mesoscale volumetric imaging of neural activity with synaptic resolution. Nat. Methods 17, 291–294 (2020).

30. W. Amir, R. Carriles, E. E. Hoover, T. A. Planchon, C. G. Durfee, J. A. Squier, Simultaneous imaging of multiple focal planes using a two-photon scanning microscope. Opt. Lett. 32, 1731–1733 (2007)

31. A. Cheng, J. Gonçalves, P. Golshani, K. Arisaka, C. Portera-Cailliau, Simultaneous two-photon calcium imaging at different depths with spatiotemporal multiplexing. Nat. Methods 8, 139–142 (2011).

32. D. Tsyboulski, N. Orlova, F. Griffin, S. Seid, J. Lecoq, P. Saggau. https://doi.org/10.1101/503052(2018).

33. J. Wu, Y. Liang, S. Chen, et al. Kilohertz two-photon fluorescence microscopy imaging of neural activity in vivo. Nat Methods 17, 287–290 (2020).

34. D. R. Beaulieu, I. G. Davison, K. Kılıç, T. G. Bifano, J. Mertz, Simultaneous multiplane imaging with reverberation two-photon microscopy. Nat. Methods 17, 283–286 (2020).

35. T. Zhang, O. Hernandez, R. Chrapkiewicz, et al., Kilohertz two-photon brain imaging in awake mice. Nat Methods 16, 1119–1122 (2019).

36. Y.-H. Tsai, C.-W. Liu, W.-K. Lin, et al. https://www.biorxiv.org/content/10.1101/2020.10.21.349712v1.full (2020).

37. T. L. Daigle, L. Madisen, T. A. Hage, et al., A Suite of Transgenic Driver and Reporter Mouse Lines with Enhanced Brain-Cell-Type Targeting and Functionality. Cell 174, 465–480 (2018).

38. N. Horton, K. Wang, D. Kobat, D. et al. In vivo three-photon microscopy of subcortical structures within an intact mouse brain. Nature Photon 7, 205–209 (2013).

39. B. G. Cooper, T. F. Manka, S. J. Y. Mizumori, Finding your way in the dark: the retrosplenial cortex contributes to spatial memory and navigation without visual cues. Behav. Neurosci. 115, 1012–1028 (2001).

40. C. Stringer, M. Pachitariu, N. Steinmetz, et al., Spontaneous behaviors drive multidimensional, brainwide activity. Science 364 (2019).

41. H. Dana, Y. Sun, B. Mohar, et al., High-performance calcium sensors for imaging activity in neuronal populations and microcompartments. Nat. Methods 16, 649–657 (2019).

42. M. Rigotti, O. Barak, M. Warden, M., X.-J. Wang, N. D. Daw, E. K. Miller, S. Fusi, The importance of mixed selectivity in complex cognitive tasks. Nature 497, 585–590 (2013).

43. L. F. Abbott, P. Dayan, The Effect of Correlated Variability on the Accuracy of a Population Code. Neural Comput. 11, 91–101 (1999). NC

44. N. A. Steinmetz, P. Zatka-Haas, M. Carandini, K. D. Harris, Distributed coding of choice, action and engagement across the mouse brain. Nature 576, 266–273 (2019).

45. C. Runyan, E. Piasini, S. Panzeri, C. D. Harvey, Distinct timescales of population coding across cortex. Nature 548, 92–96 (2017).

46. N. Li, K. Daie, K. Svoboda, S. Druckmann, Robust neuronal dynamics in premotor cortex during motor planning. Nature 532, 459–464 (2016).

47. M. W. Howe, D. A. Dombeck, Rapid signalling in distinct dopaminergic axons during locomotion and reward. Nature 535, 505–510 (2016).

48. P. Rajasethupathy, S. Sankaran, J. H. Marshel, C. K. Kim, E. Ferenczi, S. Y. Lee, A. Berndt, C. Ramakrishnan, A. Jaffe, M. Lo, C. Liston, K. Deisseroth, Projections from neocortex mediate top-down control of memory retrieval. Nature 526, 653–659 (2015).

49. E. A. Pnevmatikakis, A. Giovannucci, NoRMCorre: An online algorithm for piecewise rigid motion correction of calcium imaging data. J. Neuroscience Meth. 291, 83–94 (2016).

50. E. A. Pnevmatikakis, D. Soudry, Y. Gao, T. Machado, J. Merel, D. Pfau, T. Reardon, Y. Mu, C. Lacefield, W. Yang, M. Ahrens, R. Bruno, T. M. Jessell, D. S. Peterka, R. Yuste, L. Paninski, Simultaneous denoising, deconvolution, and demixing of calcium imaging data. Neuron 89, 285–299 (2016).

51. A. Giovannucci, J. Friedrich, P. Gunn, et al., CaImAn: an open source tool for scalable calcium imaging data analysis. eLife 38173 (2019).

52. Allen Institute for Brain Science, Allen Brain Atlas: Mouse Brain. mouse.brain-map.org (2018).

53. E. R. Kandel, J. H. Schwartz, T. M. Jessell, Principles of neural science (McGraw-Hill, Health Professions Division, New York, 2000).

54. A. Mathis, P. Mamidanna, K. M. Cury, T. Abe, V. N. Murthy, M. Weygandt Mathis, M. Bethge, DeepLabCut: markerless pose estimation of user-defined body parts with deep learning. Nat. Neurosci. 21, 1281–1289 (2018).

55. T. Nath, A. Mathis, A. C. Chen, A. Patel, M. Bethge, M. Weygandt Mathis, Using DeepLabCut for 3D markerless pose estimation across species and behaviors. Nat. Protoc. 14, 2152–2176 (2019).

56. J. M. Stujenske, T. Spellman, J. A. Gordon, Modeling the Spatiotemporal Dynamics of Light and Heat Propagation for In Vivo Optogenetics, Cell Rep. 12, 525–534 (2015).

57. K. Podgorski, G. Ranganathan, Brain heating induced by near-infrared lasers during multiphoton microscopy, J. Neurophysiol. 116, 1012–1023 (2016).

