## Supplementary Information for "High-Speed, Cortex-Wide Volumetric Recording of Neuroactivity at Cellular Resolution using Light Beads Microscopy"

**Contents:**

**Supplementary figures and captions**

**Supplementary movie captions**

**Supplementary note 1: Spatiotemporal efficiency maximization strategies**

**Supplementary note 2: MAxiMuM post objective calibration procedure and results**

**Supplementary note 3: Derivation of optimized sampling conditions for somatic imaging**

### Supplementary Figures and Captions

**Fig. S1**

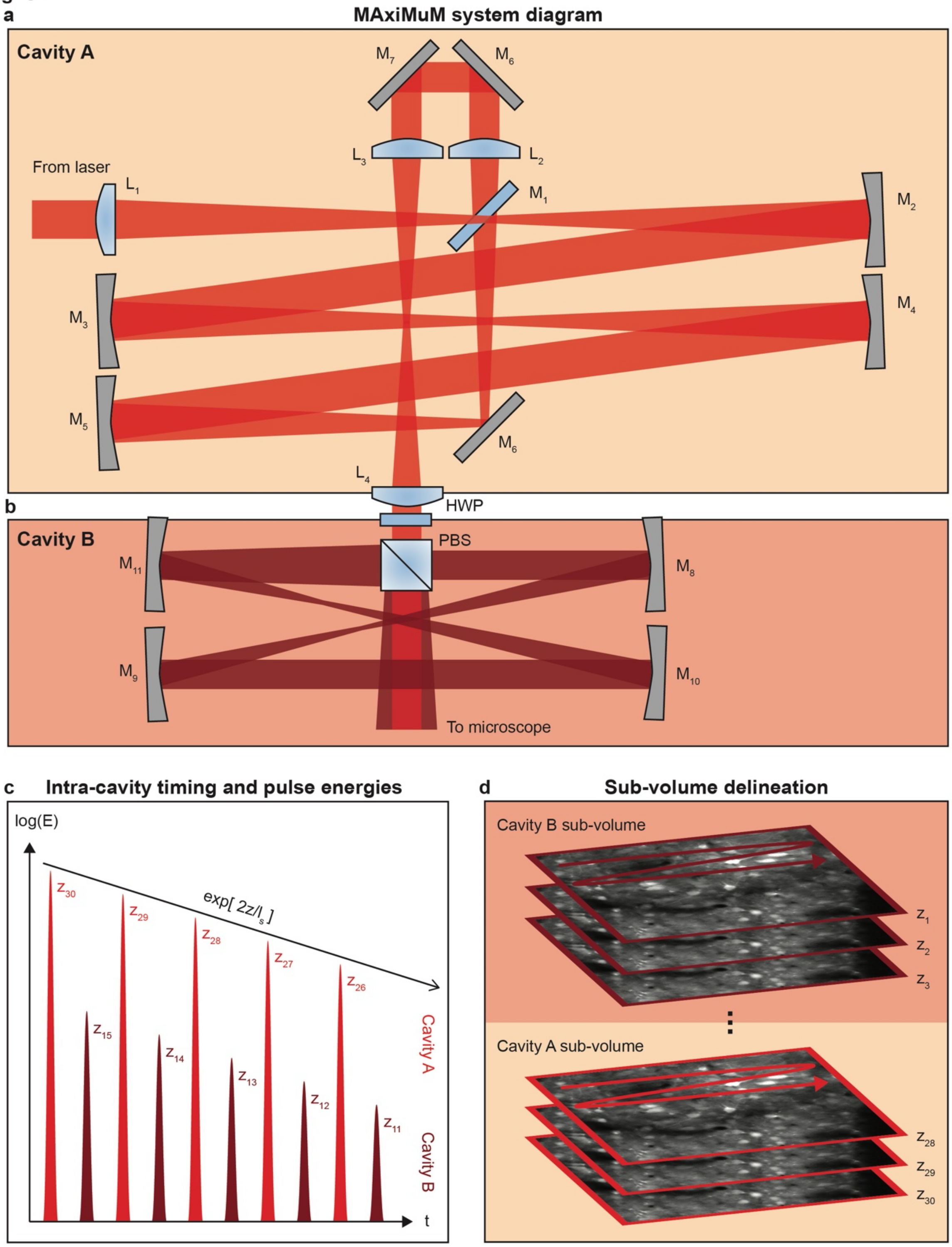

**Fig. S1: Full schematic of MAxiMuM. a,** Primary cavity schematic, where 'Ms' denote mirrors, 'Ls' denote lenses, and 'HWP' denotes a half-wave plate (see main text and Methods for design concept and description of beam propagation through the system). **b,** A second cavity with shorter focal length mirrors creates a copy of the 15 pulses from Cavity A and shifts them in time and axial location to achieve the full 30 beams and 465 μm axial range of the MAxiMuM system. **c,** Temporal schematic of pulses from cavity A and B. Due to the shorter delay of cavity B relative to A, the pulse trains are interleaved. The pulse energies for each beam decrease exponentially due to the partially transmissive mirror ($M_1$) in cavity A. Exponential decrease is matched to the expected scattering length ($l_s$) for brain tissue (~200 μm). Power for cavity B pulses are lower than those from A since cavity B pulses are sent to more superficial layers in the brain; relative power can be controlled by the HWP in **a**. **d,** Sub-volume schematic. Cavities A and B form two sub-volumes, with the planes from cavity A below those from B such that together they continuously sample the entire axial range.

Fig. S2

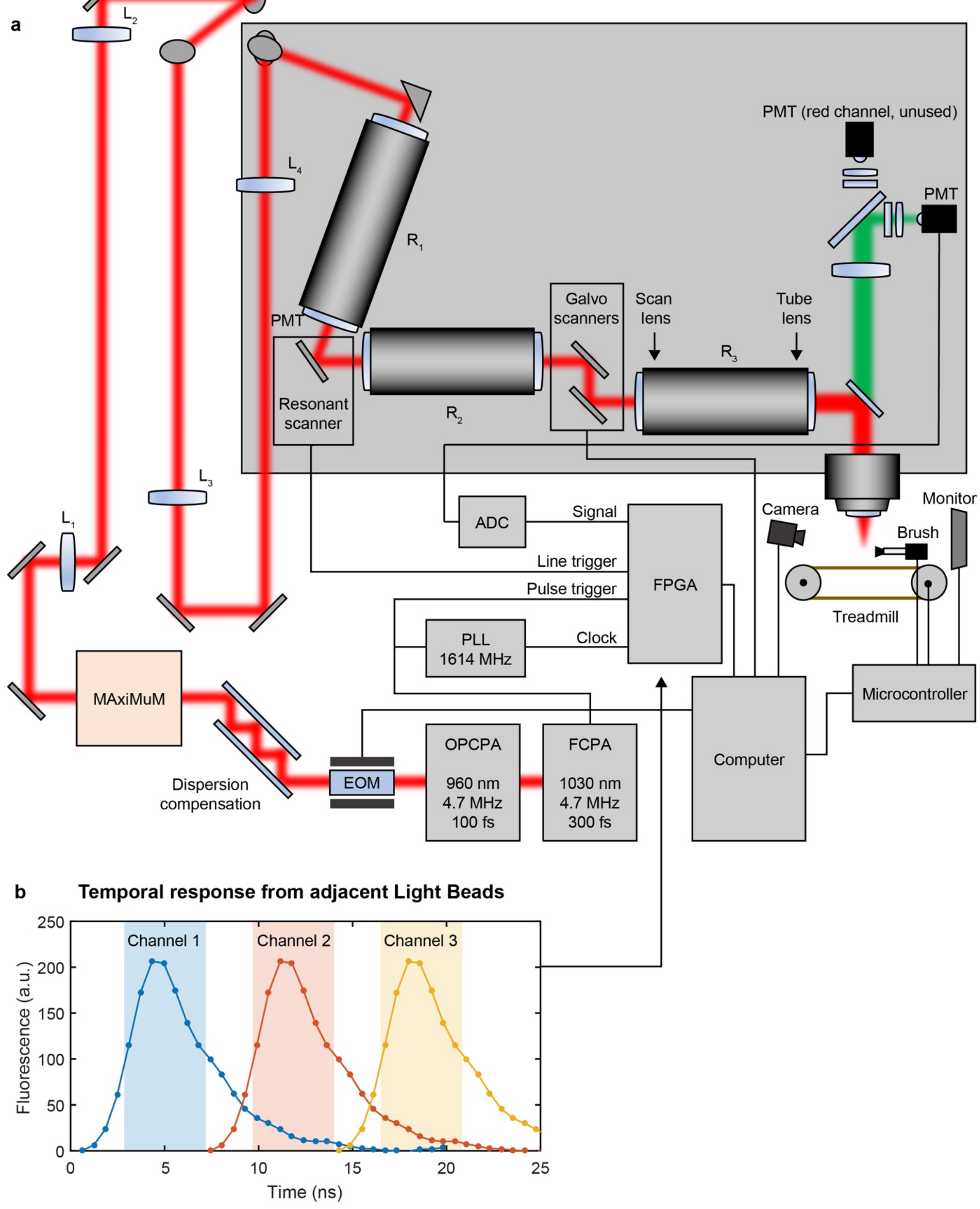
a
Overview of Light Beads Microscopy Setup
L2
L4
L3
L1
R1
R2
R3
PMT (red channel, unused)
PMT
PMT
Galvo scanners
Scan lens
Tube lens
Resonant scanner
ADC
Signal
Line trigger
Pulse trigger
FPGA
PLL 1614 MHz
Clock
Camera
Monitor
Brush
Treadmill
Microcontroller
Computer
MAxiMuM
Dispersion compensation
EOM
OPCPA
960 nm
4.7 MHz
100 fs
FCPA
1030 nm
4.7 MHz
300 fs
b
Temporal response from adjacent Light Beads
Channel 1
Channel 2
Channel 3
Fluorescence (a.u.)
Time (ns)
0
50
100
150
200
250
0
5
10
15
20
25

**Fig. S2: Full microscope schematic. a,** Setup schematic for mesoscope system starting from the fiber chirped-pulse amplifier (FCPA), through the optical parametric chirped-pulse amplifier (OPCPA), electro-optic modulator (EOM), dispersion compensation path, MAxiMuM, and into the microscope. 'Ls' denote lenses, 'Rs' denote relay lens pairs, 'PMT' denotes photo-multiplier tube, 'ADC' denotes analog to digital converter, and 'PLL' denotes phase-locked loop. **b,** Schematic showing channel allocation for demultiplexing of signal from three adjacent light beads on the FPGA. Data points are the measured impulse response for fluorescence from GCaMP6f measured with our PMT and associated electronics, captured with 1614 MHz (0.62 ns) resolution. Shaded regions denote the integration boundaries for each de-multiplexed channel.

Fig. S3

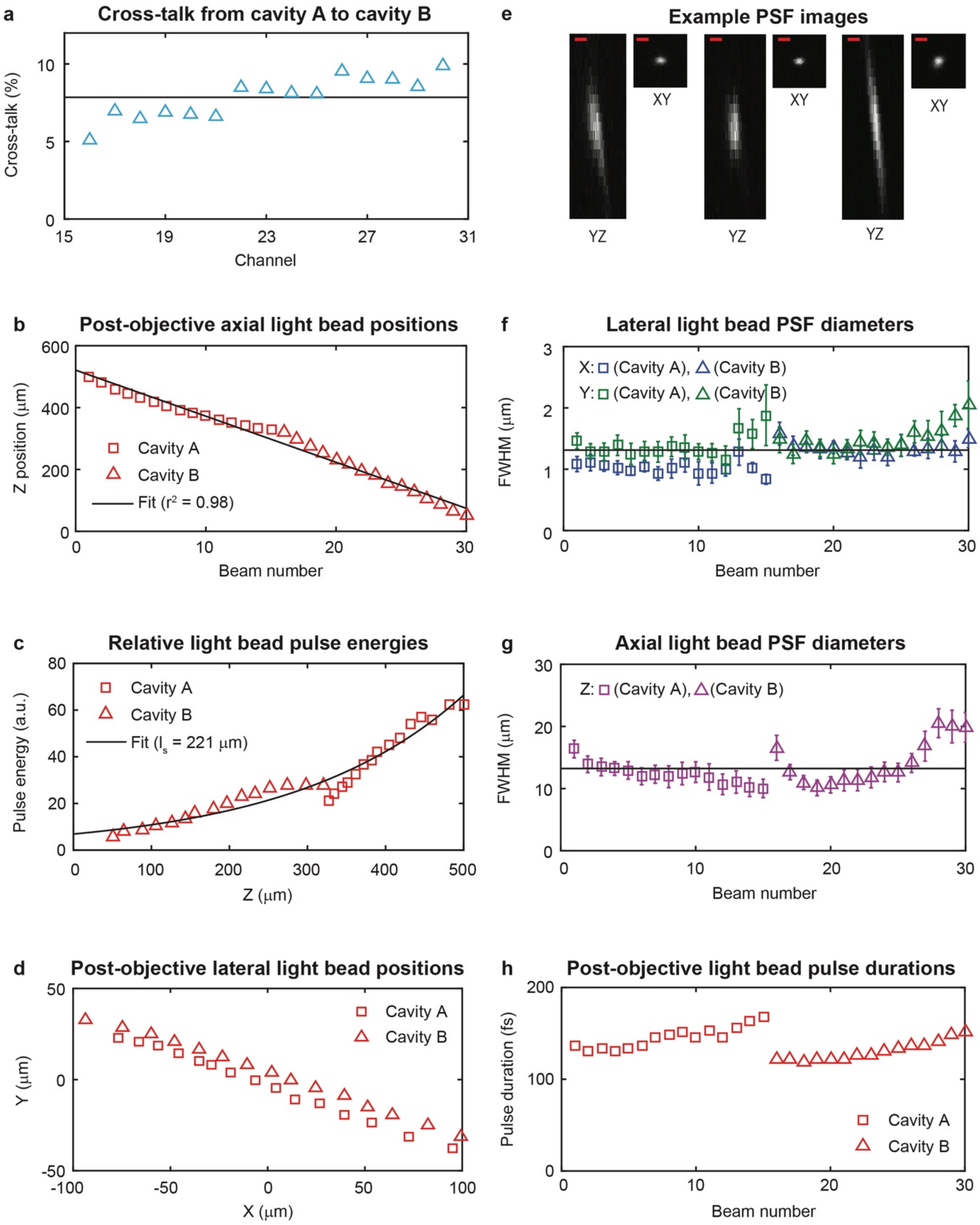
a
Cross-talk from cavity A to cavity B
Cross-talk (%)
Channel
e
Example PSF images
XY
YZ
b
Post-objective axial light bead positions
Z position (µm)
Cavity A
Cavity B
Fit (r² = 0.98)
Beam number
f
Lateral light bead PSF diameters
X: (Cavity A), (Cavity B)
Y: (Cavity A), (Cavity B)
FWHM (µm)
Beam number
c
Relative light bead pulse energies
Pulse energy (a.u.)
Cavity A
Cavity B
Fit (ls = 221 µm)
Z (µm)
g
Axial light bead PSF diameters
Z: (Cavity A), (Cavity B)
FWHM (µm)
Beam number
d
Post-objective lateral light bead positions
Y (µm)
Cavity A
Cavity B
X (µm)
h
Post-objective light bead pulse durations
Pulse duration (fs)
Cavity A
Cavity B
Beam number

**Fig. S3: Post-objective calibration of light bead columns generated by MAxiMuM. a,** Measurement of crosstalk between demultiplexed channels. Black horizontal line shows mean value at ~7%. **b–d,** Axial position, relative pulse energy, and transverse position of the light beads from MAxiMuM, respectively, calibrated by translating a pollen grain through the focus of the microscope. **e-g**, characterization of light bead point spread functions. **e,** example images of PSFs for light beads 5, 15, and 25 in the *x,y* and *x,z* planes. Scale bar: 2 μm. **f,** Lateral point-spread function full-width at half-maximum diameters for each light bead. Mean value shown by horizontal black line. Error bars denote the 95% confidence interval values for the Gaussian fits used to determine PSF widths. **g,** Axial point-spread function full-width at half-maximum diameters for each light bead. Mean value shown by horizontal black line. Error bars denote the 95% confidence interval values for the Lorentzian fits used to determine PSF widths. **h,** Pulse duration measurements of each beam from MAxiMuM, post-objective.

**Fig. S4** **Schematics of data processing pipeline**

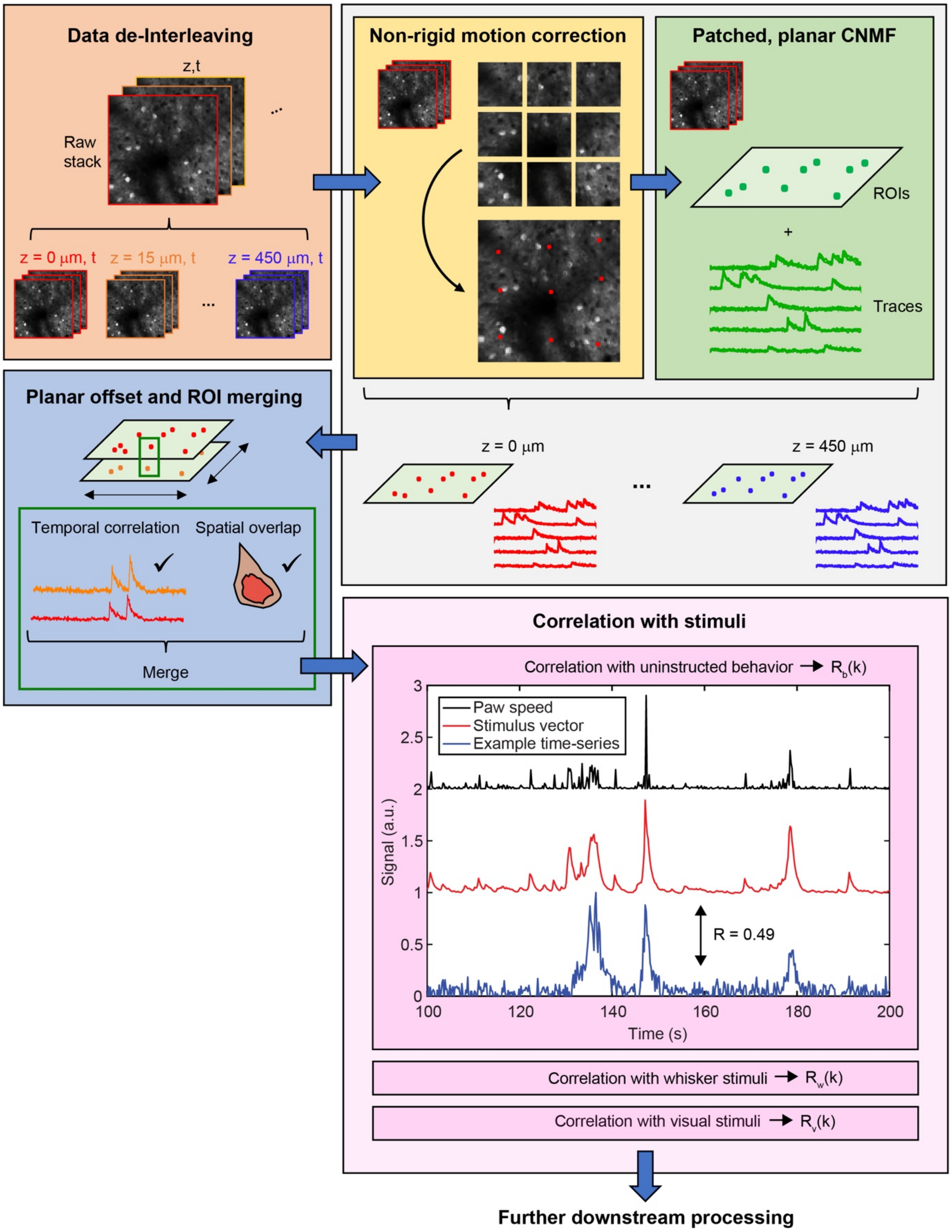

**Fig. S4: Schematic of the data processing pipeline**. Raw data is assembled into frames and separated into individual temporal stacks for each *z* plane. Each stack is separately motion-corrected and sent through a constrained non-negative matrix factorization (CNMF) sub-routine to extract neuronal footprints and time-series. Lateral offsets between the planes are accounted for using calibration values, and neurons with both correlated temporal activity and overlapping spatial footprints are merged to prevent doubly-counted cells. Neurons are correlated with vectors representing each stimulus to determine if they are tuned. Example raw and kernel-convolved stimulus vectors and an example time-series for a tuned neuron are shown for uninstructed behaviors during a recording.

**Fig. S5**

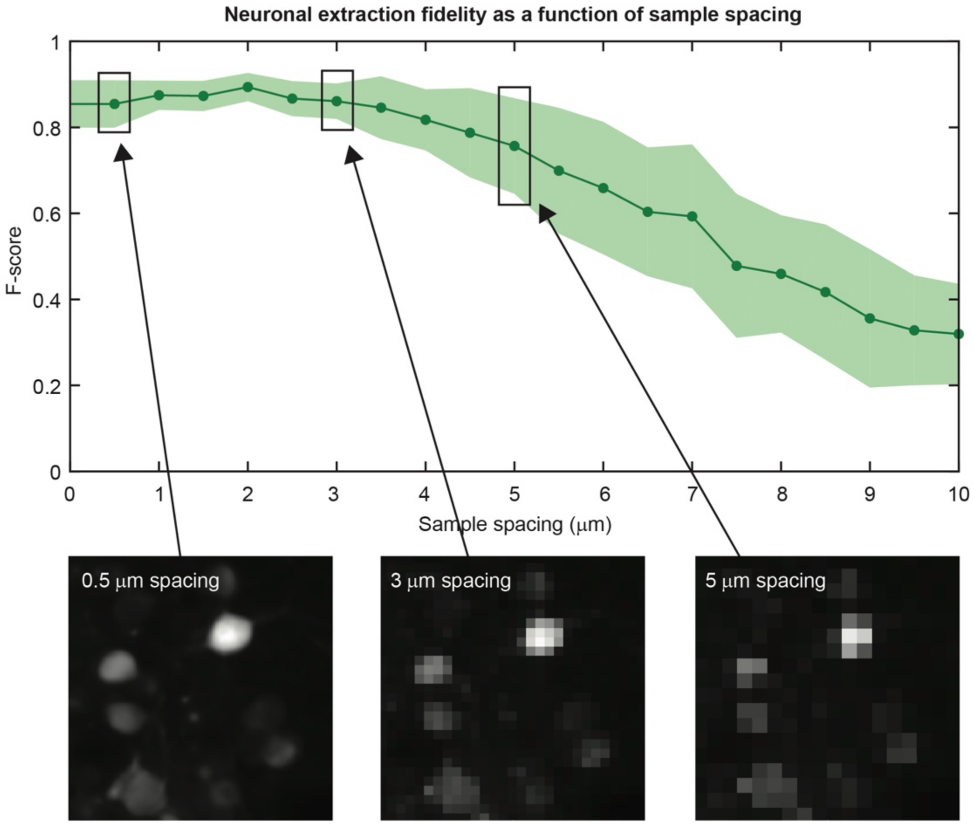
Neuronal extraction fidelity as a function of sample spacing
F-score
Sample spacing (μm)
0
0.2
0.4
0.6
0.8
1
0
1
2
3
4
5
6
7
8
9
10
0.5 μm spacing
3 μm spacing
5 μm spacing

**Fig. S5. Extraction fidelity, measured by F-score, as a function of sample spacing.** F-score is defined as the harmonic mean of the sensitivity and precision of the neuronal extraction. Solid line indicates mean value, shaded region indicates one standard deviation from the mean. Example images for 0.5, 3, and 5 μm sample spacing inset.

**Fig. S6**

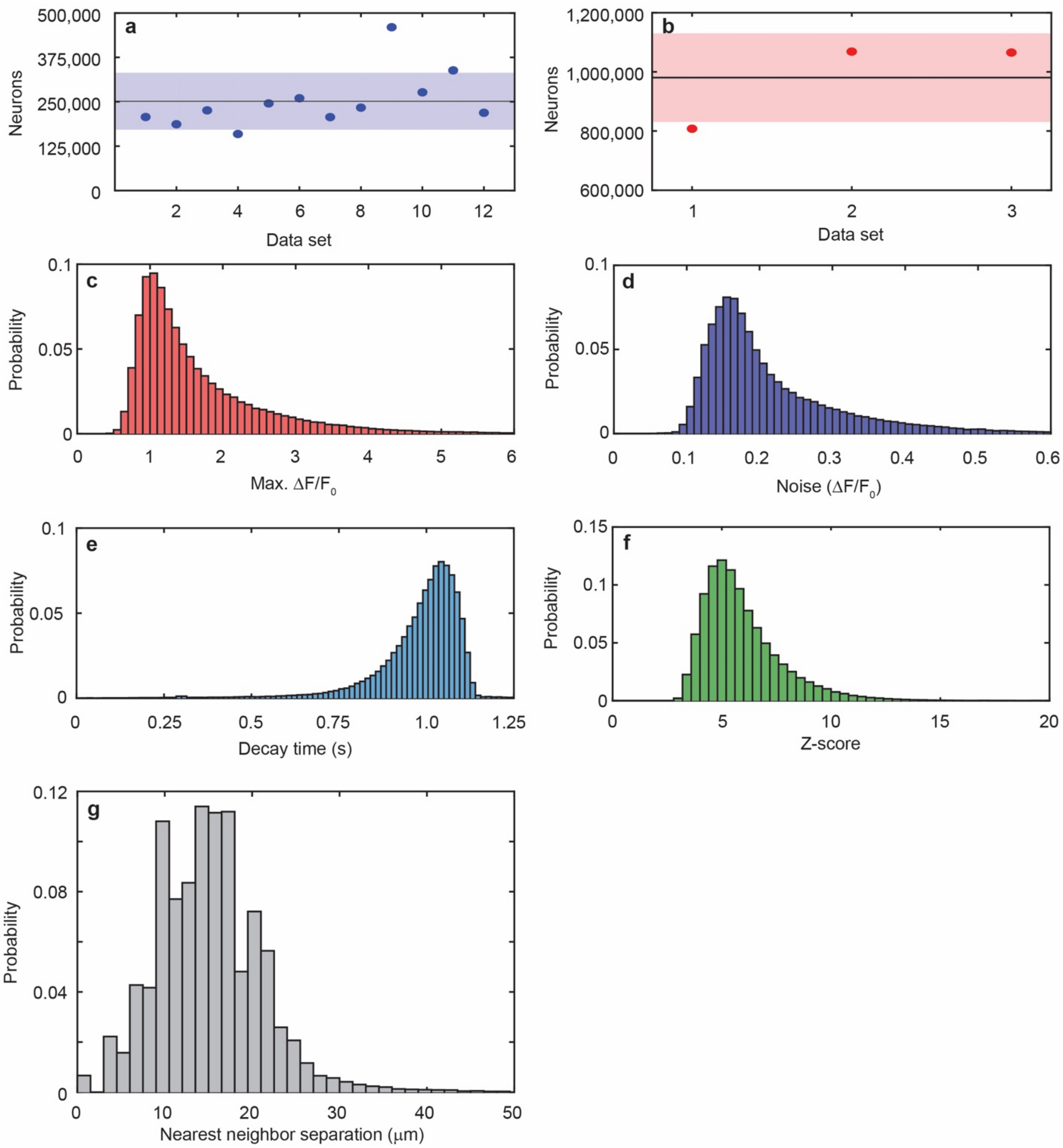

a
500,000
375,000
250,000
125,000
0
Neurons
2
4
6
8
10
12
Data set
b
1,200,000
1,000,000
800,000
600,000
Neurons
1
2
3
Data set
c
0.1
0.05
0
Probability
0
1
2
3
4
5
6
Max. ΔF/F0
d
0.1
0.05
0
Probability
0
0.1
0.2
0.3
0.4
0.5
0.6
Noise (ΔF/F0)
e
0.1
0.05
0
Probability
0
0.25
0.5
0.75
1.0
1.25
Decay time (s)
f
0.15
0.1
0.05
0
Probability
0
5
10
15
20
Z-score
g
0.12
0.08
0.04
0
Probability
0
10
20
30
40
50
Nearest neighbor separation (μm)

**Fig. S6 Indicator and extraction statistics. a,** Summary of the number of neurons extracted from 12 recordings with ~3 × 5 × 0.5 $mm^3$ FOV across N = 6 animals; solid black line denotes the mean, shaded region denotes 1 standard deviation from the mean; data set 1 corresponds to the recording shown in Figs. 2 and 3. **b,** Summary of the number of neurons extracted from 3 recordings with ~5.4 × 6 × 0.5 $mm^3$ FOV across N = 3 animals; solid black line denotes the mean, shaded region denotes 1 standard deviation from the mean; data set 1 corresponds to the recording shown in Fig. 5. **c,** Distribution of maximum $\Delta F/F_0$ values for transients measured in mice expressing GCaMP6s from the experiment shown in Fig. 2. **d,** Distribution of baseline noise levels (expressed in terms of $\Delta F/F_0$) of the population in **c**. **e,** Distribution the average transient decay time for the traces in the population in c. **f,** Distribution of maximum Z-score for each neuron from the population shown in Fig. 2 with SNR threshold of 1.4. **g,** Nearest neighbor distances for the population of neurons shown in Fig. 2.

**Fig. S7**

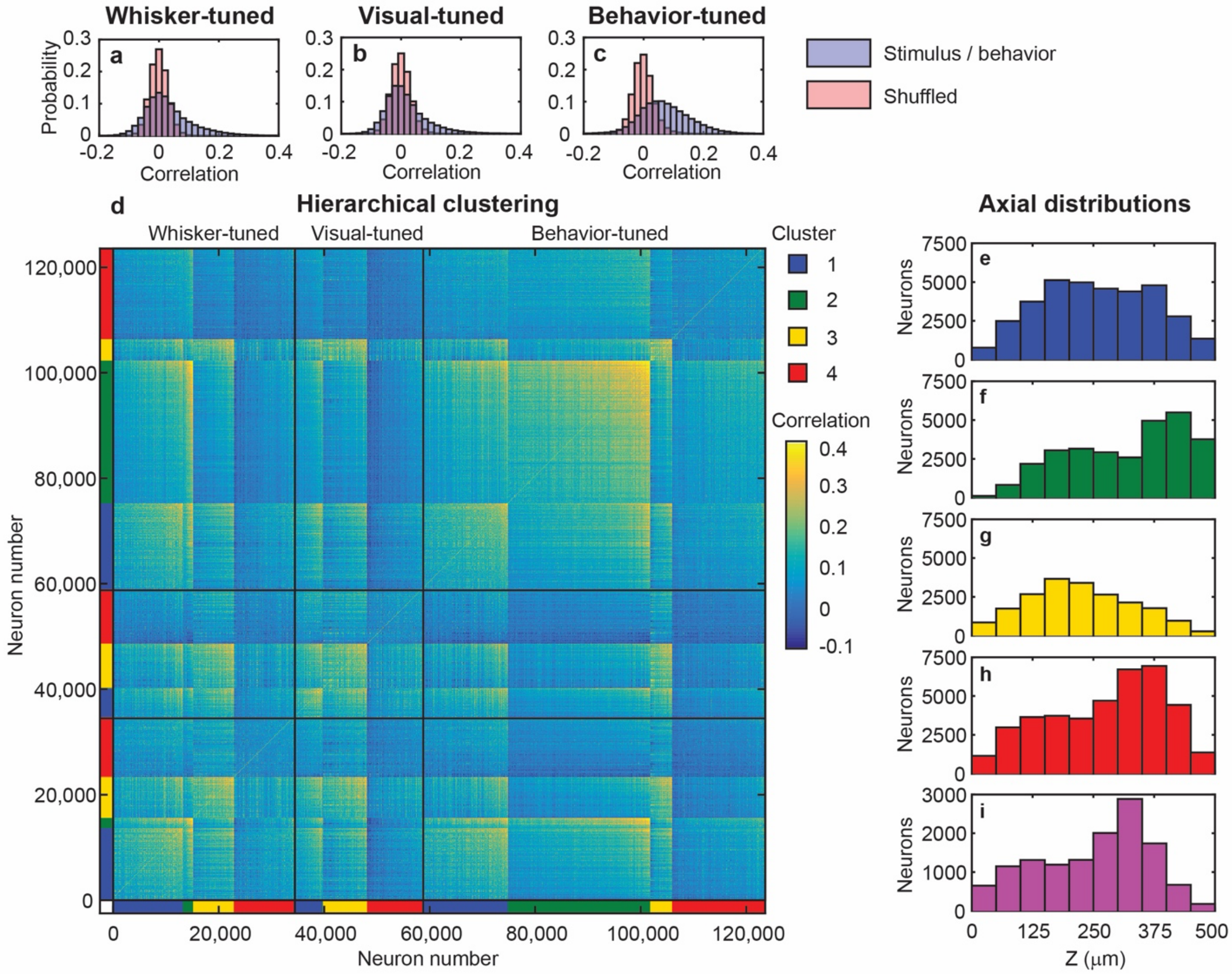
Whisker-tuned
Visual-tuned
Behavior-tuned
a
b
c
Probability
Correlation
Stimulus / behavior
Shuffled
d
Hierarchical clustering
Whisker-tuned
Visual-tuned
Behavior-tuned
Cluster
1
2
3
4
Correlation
Neuron number
Neuron number
Axial distributions
e
f
g
h
i
Neurons
Z (μm)

**Fig. S7: Clustering analysis of stimulus-tuned neurons from the dataset shown in Fig. 2. a–c**, Correlation distributions of neurons with whisker stimuli, visual stimuli, and uninstructed spontaneous animal behaviors (blue), compared to time-shuffled distributions (red). **d,** Correlation matrix of all neurons tuned to any stimulus condition. The matrix is sorted by stimulus (boundaries denoted by black lines), cluster, and mean correlation. **e–i,** Axial spatial distributions of neurons in clusters 1 through 4 and the uncorrelated population, respectively.

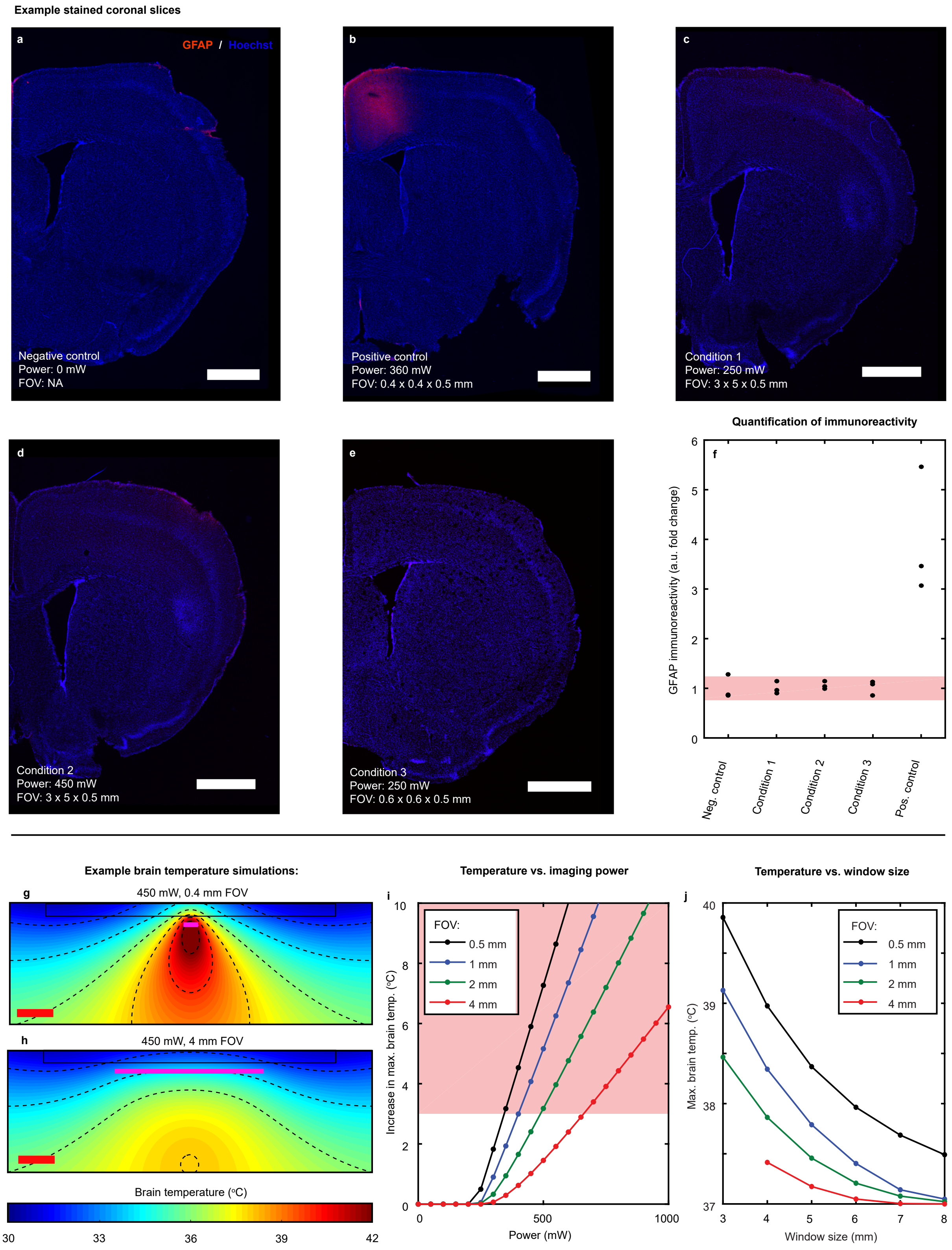
Example stained coronal slices
a
GFAP / Hoechst
Negative control
Power: 0 mW
FOV: NA
b
Positive control
Power: 360 mW
FOV: 0.4 x 0.4 x 0.5 mm
c
Condition 1
Power: 250 mW
FOV: 3 x 5 x 0.5 mm
d
Condition 2
Power: 450 mW
FOV: 3 x 5 x 0.5 mm
e
Condition 3
Power: 250 mW
FOV: 0.6 x 0.6 x 0.5 mm
Quantification of immunoreactivity
f
GFAP immunoreactivity (a.u. fold change)
0
1
2
3
4
5
6
Neg. control
Condition 1
Condition 2
Condition 3
Pos. control
Example brain temperature simulations:
g
450 mW, 0.4 mm FOV
h
450 mW, 4 mm FOV
Brain temperature (°C)
30
33
36
39
42
Temperature vs. imaging power
i
FOV:
0.5 mm
1 mm
2 mm
4 mm
Increase in max. brain temp. (°C)
0
2
4
6
8
10
0
500
1000
Power (mW)
Temperature vs. window size
j
FOV:
0.5 mm
1 mm
2 mm
4 mm
Max. brain temp. (°C)
37
38
39
40
3
4
5
6
7
8
Window size (mm)

**Figure S8: Brain heating experimental data and simulations. a-e,** Representative images of brain sections showing immunolabeling for astrocyte activation marker (anti-GFAP, red) and DNA stain (Hoechst 33342, blue) after exposure to the laser power and FOV listed below. Scale bars: 1 mm **a,** control, no laser exposure. **b,** 360 mW, 0.4×0.4×0.5 mm FOV (2250 $mW/mm^2$). **c,** 250 mW, 3×5×0.5 mm FOV (17 $mW/mm^2$). **d,** 450 mW, 3×5×0.5 mm FOV (34 $mW/mm^2$). **e,** 250 mW, 0.6×0.6×0.5 mm FOV (700 $mW/mm^2$). **f,** Intensity of immunolabeling corresponding to imaging intensity as a fraction compared to mean of control samples. $N$ = 3 separate brain hemispheres per condition. Shaded area denotes the 95% confidence interval of the control group mean. **g,h,** Simulations of brain temperature at steady state for 450 mW of optical power in a (**g**) 0.4 mm FOV and a (**h**) 4 mm FOV; magenta lines indicated nominal focal plane; scale bars: 1 mm. Brain temperature in **g** heats to ~6 ºC above core temperature (37 ºC), while heating in **h** is only ~1 ºC. Solid black lines denote boundaries of the cranial window, magenta lines indicate focal plane, dashed black lines indicate contours separated by 2 ºC. Scale bar: 1 mm. **i,** Maximum brain temperature as a function of optical power and FOV. Cooling of the brain through the cranial window leads to a minimum power threshold before the onset of heating in the brain. **j,** Brain temperature for a fixed power level (250 mW) as a function of cranial window diameter and FOV. Larger cranial windows lead to less overall heating of the brain.

### Supplementary movie captions

**Movie S1: Example mouse behavior during recording.** Motion tracking of the right and left paws and right ankle are shown with green, blue, and red dots respectively.

**Movie S2: 3D rendering of the top 56,954 most active neurons from 207,030 detected neurons within a recording volume of ~3 × 5 × 0.5 mm recorded at 4.7 Hz from the data set shown in Fig. 2.** Occurrences of whisker stimuli, visual stimuli, or simultaneous whisker and visual stimuli are denoted by the red mark in the upper left-hand corner of the frame. Playback sped up 4×.

**Movie S3: Example recording of a single plane (depth = 344 μm) from the volumetric recording shown in Fig. 2 (3 × 5 mm FOV recorded at 4.7 Hz).** Playback sped up 4×. Scale bar: 250 μm.

**Movie S4: Summary of all traces extracted from the recording in Fig. 2.**

**Movie S5: Time-lapse of activity onset for neurons tuned to uninstructed behaviors of the animal following the analysis shown in Figs. 3r and 3s.** Playback is displayed in real time.

**Movie S6: Example recording of a single plane (depth = 384 μm) from the volumetric recording shown in Figs. 4a–4c (0.6 × 0.6 mm FOV recorded at 9.6 Hz).** Playback sped up 4×. Scale bar: 50 μm.

**Movie S7: 3D rendering of the top 32,915 most active neurons from 70,275 detected neurons within a recording volume of ~2 × 2 × 0.5 mm recorded at 6.5 Hz from the data set shown in Figs. 4d–4g.** Occurrences of whisker stimuli are denoted by the red mark in the upper left-hand corner of the frame. Playback sped up 4×.

**Movie S8: Example recording of a single plane (depth = 144 μm) from the volumetric recording shown in Figs. 4d–4g (2 × 2 mm FOV recorded at 6.5 Hz).** Playback sped up 4×. Scale bar: 200 μm.

**Movie S9: 3D rendering of the top 150,000 most active neurons from 807,748 detected neurons within a recording volume of ~5.4 × 6 × 0.5 mm recorded at 2.2 Hz from the data set shown in Fig. 5.** Playback sped up 4×.

**Movie S10: Example recording of a single plane (depth = 600 μm) from the volumetric recording shown in Fig. 5 (5.4 × 6 mm FOV recorded at 2.2 Hz).** Playback sped up 4×. Scale bar: 500 μm.

**Movie S11: Summary of all traces extracted from the recording in Fig. 5.**

**Supplementary Notes**

**Supplementary Note 1**

The following note outlines crucial trade-offs for designing maximally-efficient sampling for mesoscopic scan engines and highly-multiplexed systems. Each subsection shows the relationships between different coupled parameters in the design.

*Resolution, repetition rate, and resonant scanner*

Design choices for the laser repetition rate and resonant scanner speed are coupled to the desired resolution in the sample. Given a desired lateral pixel resolution $\delta x$, and the lateral extent of the resonant scan path $X_{res}$, we can determine the desired number of pixels $N_x$:

$$N_x = \frac{X_{res}}{\eta\ \delta x}$$

Because the resonant scanner has a sinusoidal trajectory, as it reverses direction there will be an inevitable number of pixels during this turn-around time that do not contribute to the image; the fill fraction $\eta$ accounts for this loss in the equation above.

In the one pulse per pixel regime, the number of pixels required drives the necessary repetition rate of the laser $f_{laser}$ and the frequency of the resonant scanner $f_{resonant}$ such that $N_x = \frac{f_{laser}}{2\ f_{resonant}}$, where the factor of 2 assumes bi-directional scanning. Thus, the necessary laser repetition rate is given by:

$$f_{laser} = \frac{2\ f_{resonant}\ X_{res}}{\eta\ \delta x}$$

For our system, we employ a $f_{resonant}$ = 12 kHz resonant scanner with a lateral extent of $X_{res}$ = 600 µm. Therefore, for a desired resolution of $\delta x$ = 5 µm and a typical fill fraction of $\eta$ = 0.75, the optimum laser repetition rate is 3.8 MHz. Our laser is slightly faster (4.7 MHz), hence the increase in our lateral spatial resolution along the resonant scan dimension (4.2 µm).

*Field of view (FOV) and frame rate*

We can explore tradeoffs between the FOV and frame rate of the microscope based on the choice of resonant scanner and desired pixel resolution. In a multiple-galvo and resonant scanner scanning configuration, the resonant scanner and a galvo mirror scanning the orthogonal (*y*) direction form a rectangular region of interest that can be tiled an integer number $n_x$ times in the *x* direction to increase the FOV. The lateral extent and spatial sampling in the *y* direction, *Y* and $\delta y$ respectively, can be tailored independent of the *x* direction (which is necessarily coupled to the resonant scanner). Thus, the FOV is given by $FOV = n_x X_{res} Y$ and we can estimate the frame rate $f_{frame}$ as:

$$f_{frame} = 2f_{resonant} \frac{\delta y}{n_x Y}$$

For the 3 × 5 mm FOVs demonstrated here (Figs. 2 and 3), $n_x$ = 5 regions are tiled together, each with dimensions $X_{res} \times Y$ = 0.6 × 5 mm, and $\delta y$ = 5 µm pixel spacing in the *y* direction, yielding an estimated frame rate of 4.8 Hz. Note that the flyback times of the galvos in the system add some overhead depending on the scan field and the speed of the scanners, however, the reduction in speed is negligible to first order.

*Multiplexing design*

To maximize temporal efficiency, the system should operate as close to the maximum voxel rate $f_{voxel}$ possible given the limit imposed by the fluorescent lifetime of the indicator (for GCaMP, $f_{voxel} \sim 150$ MHz). The laser repetition rate and maximum voxel rate thus determine the upper limit of the degree of multiplexing $N$:

$$N = \left\lfloor \frac{f_{voxel}}{f_{laser}} \right\rfloor$$

For our case, this works out to a maximum degree of multiplexing $N = 31$, however, the use of the second cavity necessitated an even number of beams, hence our $N = 30$ operation point.

It should be noted that the maximum voxel rate is a function of the fluorescence lifetime *convolved* with the speed of the detection mechanism, as well as by the tolerable crosstalk between beams. While we employed a typical photomultiplier tube for our experiments, hybrid photomultiplier tubes could potentially increase the speed of measurement, and thus increase maximum possible $N$, at the cost of reduced gain and lower signal levels. Our experiments were also very conservative with respect to crosstalk – one could increase $N$ by spacing beams closer together in time, inviting more crosstalk. The majority of the detrimental effects could be removed in post-processing through *a priori* calibration of the mean crosstalk between bins. This approach is very successful provided the data have excellent SNR, as only the mean crosstalk induced signal, and not cross-talk induced noise, can be removed.[34]

Equation (1) in the manuscript, shows the relationship between the desired axial sampling resolution $\delta z$, the focal offset $\Delta z$, the lateral magnification of the microscope $M$, and the transmission of the cavity output coupler $T$. Assuming an 8-f configuration for the cavity, one can work out the necessary focal lengths of the concave mirrors, $f_{mirror}$, in order to achieve $N$-fold multiplexing within the repetition rate of the laser:

$$f_{mirror} = \frac{1}{4}\left[\frac{c}{N f_{laser}} - \Delta z\right]$$

where $c$ is the speed of light. For our system, the optimum works out to be $f_{mirror}$ ~ 500 mm in the main cavity (15× multiplexing), and $f_{mirror}$ ~ 250 mm in the second cavity.

*Final considerations*

By adjusting the coupled parameters described above, one can generate a prescription for the most efficient scanning engine possible prioritizing the specifications most salient for the desired application. For example, our demonstration of LBM prioritized the axial extent of the volume, $N \cdot \delta z$, and employed the minimum required lateral sampling. As an alternative, one could prioritize high spatial resolution (small $\delta x$ and $\delta y$) without sacrificing FOV by increasing the laser repetition rate — at the cost of lower possible $N$. By iteratively tuning parameters, these tradeoffs should become clear and the scan engine design will converge.

**Supplementary Note 2**

The following note details the methods and results for post-objective calibration of MAxiMuM including crosstalk analysis, as well as characterization of the power, position, PSF, and pulse duration of each light bead.

A crucial element of the spatiotemporal multiplexing scheme employed here is that signal from the beams is distinguishable in time in order to effectively de-multiplex and assign signal to the proper axial location. Any leakage of the signal from one beam to the time slot of another will lead to distortion of the volume, typically referred to as crosstalk. We measured crosstalk in our system by recording mouse brain tissue *in vivo* with 30× multiplexing enabled, but with the beams in cavity B (Fig. S1b) blocked. As shown in the channel plots inset in Fig. S2b, the fluorescence response from a single voxel has an exponential tail that inevitably extends to some degree into the signal from the subsequent voxel. The fraction of residual signal generated by the beams from cavity A, recorded at time values allocated for beams from cavity B, indicates the crosstalk for the system. As shown in Fig. S3a, the crosstalk is fairly channel-invariant and the average value (7%) is largely negligible. Further, as shown in Refs. 10, 31, and 34, crosstalk is a linear mixing phenomenon and can be removed post-recording if the contributions from each channel are known *a priori*.

Figure S3b shows a representative calibration of the relative axial focal position for each beam post-objective, achieved by recording fluorescence signal from a grain of pollen (~20 μm diameter) translated through the focus of the objective by a piezo stage (NPoint, NPFocus1000). The offset between axial foci of the beams is linear ($r^2$ = 0.98) over the 30 beams. Additionally, by tracking the maximum time-averaged signal from a pollen grain measured at the nominal focal plane of each beam, we could calibrate the power in each plane (Fig. S3c). We fit the power measurements with an exponential curve, finding that the effective scattering length ($l_s$ = 220 μm) matched the expectation for the sample ($l_s$ ~ 200 μm) reasonably well. Pollen grain measurements were also used to determine the lateral offset between the beams (Fig. S3d) which corresponded to less than 200 μm total shift across the axial range.

The point spread function (PSF) for each beam was also recorded using a 1 μm diameter fluorescent bead (Tetraspeck T14792). Example X-Z projections of the PSF of several light beads is shown in Fig. S3e, and the measured full-width-at-half-maximum (FWHM) diameter for the PSF of each beam is shown for the lateral and axial dimensions in Fig. S3f and S3g respectively. The measured axial widths ensure that for the given separation between axial planes (~15 μm) and typical neuron sizes (10 – 20 μm), the probability of detecting any neurons present is high, while the likelihood of the PSF intersecting two neurons simultaneously (thus rendering them indistinguishable) is negligibly low. The average lateral FWHM is 1.3 μm, which when deconvolved (assuming a step function with 1 μm width as a model for the fluorescent bead) yields an average PSF diameter of 1.1 μm.

Care was taken to minimize the inter-cavity dispersion for MAxiMuM. While dispersion accumulation throughout the microscope is inevitable and can be compensated, any dispersion within the cavity would lead to *differential* broadening of the pulse for each beam exiting MAxiMuM. Accordingly, we designed the cavity with entirely reflective components and used low-dispersion dielectric coatings for as many components as possible. Figure S3h shows the post-objective pulse durations measured by auto-correlation for each of the 30 beams (APE PulseCheck). While the pulse duration (120 – 160 fs) is not transform-limited (~90 fs) for each beam after the objective, the degradation is acceptably small and can be accounted for by adjusting the power of each beam through optimization of the output coupling from the cavity.

**Supplementary Note 3**

The following note discusses the simulation methods and analysis used to determine the optimum voxel spacing for functional imaging of neuronal somas.

A key feature of the performance of the Light Beads Microscopy approach is to optimize sampling such that spatial location *and* the time-series of neurons are faithfully extracted without sacrificing volumetric FOV. The typical repetition rate in most calcium imaging systems (~80 MHz) combined with available speeds of scanning systems (~12 kHz) and typical FOVs (~500 μm) implies a spatial oversampling of pixels such that each pixel is effectively excited by multiple pulses (~150 nm sample spacing and a typical ~500 nm PSF diameter yields >4 pulses per unique pixel). Sampling beyond the density dictated by the PSF diameter, or more generally, beyond the features of interest within the tissue, leads to inefficiency in signal generation and reduction of the volume size that can be imaged at the minimum acceptable temporal resolution. Optimizing the scanning architecture, pulse repetition rate, and FOV along with the degree of multiplexing in a top-down approach is necessary to achieve the efficient sampling required for mesoscopic and volumetric performance and has been a limitation of previous demonstrations of multiplexed two-photon microscopy.[10,19,23,25,30-34] However, sampling density cannot be arbitrarily sparse, otherwise the system would lose the capability to extract transients from single neuronal cells.

To quantitively and optimize this trade-off between efficiency and resolution, we developed a simulation pipeline. We started by recording 7 single-plane recordings of mice expressing GCaMP6f ($N = 2$) over a variety of different depths in the posterior parietal cortex. These data sets were recorded with diffraction-limited resolution (~600 nm) using the conventional operation mode of our multiphoton mesoscope with each recording having a 300 μm FOV, 0.5 μm sample spacing, and 26 Hz frame rate – consistent with a conventional two-photon microscope. We hand-segmented each data set to extract the footprints and time-series of active neurons during the recordings, finding 50 – 80 neurons in each plane. Next, we sent each data set through the motion correction and neuronal extraction portions of our data processing pipeline (see relevant section above) and compared the population and the time-series of extracted neurons to the hand-curated ground truth. We calculated the F-score of the extraction from the automated pipeline, defined as the harmonic mean of the sensitivity (true positive rate) and precision (1 – false positive rate) of the neuronal extraction, to quantify the extraction fidelity. Subsequently, we removed pixels from the data sets and re-ran the extraction to determine the F-score as a function of sampling sparsity.

Figure S5 shows the F-score as a function of the effective pixel size. The maximum F-score possible is ~0.85 due to some inevitable imperfection in the extraction algorithm. This performance is preserved up to 3 μm pixels, in keeping with expectation given that 3 μm pixel resolution is more than sufficient to sample neuronal cell bodies with typical diameters of 10 – 20 μm (see example mean projection images inset in Fig. S5). For larger pixels, the mean F-score decreases and standard deviation over all data sets analyzed (denoted by the width of the curve in Fig. S5) increases. To approximate a limit for tolerable degradation of F-score, we allow the mean F-score to fall at most one standard deviation below the average performance for the high-resolution condition. This corresponds to a pixel size of ~4.5 μm. The maximum width of each region of interest of our mesoscope combined with the resonant scanner speed and pulse repetition rate leads to a minimum pixel size of 4.2 μm in the $x$ direction. Accordingly, we increased the pixel size in the $y$ direction to 5 μm, such that the mean pixel size corresponds to the optimized value above.

**Supplementary Note 5**

The following note outlines experiments and simulations designed to determine the limits of brain safety due to heating during imaging.

The large volumes accessible by the LBM method exceed those of previously reported 2p methods by more than an order of magnitude (Table 1). However, the increased multiplicity of the method also demands an increase in the optical power used for imaging, especially when imaging deep in the brain. Naively, one expects a relationship between the intensity of the illumination and the degree of heating in the brain, as tissue absorption is an intensity-dependent process. However, it is not clear that heating scales strictly linearly with increasing intensity.[57] Therefore, we elected to validate the imaging conditions used in our experiments through immunohistochemical labeling experiments and simulations of the brain temperature. While both these approaches have been demonstrated previously in the literature for analyzing heat related-damage in multi-photon imaging, the following results expand this analysis for FOVs and absolute power levels well above what has been previously shown.

Astrocyte activation, which we have also used to validate some of our previous imaging modalities,[24] is a well-established[57] phenotype of neuropathology and neurotrauma, including those that arise due to the exposure of tissue to temperatures exceeding physiological conditions. To demonstrate that our imaging method, at all power levels used in the manuscript, does not induce any adverse effects, we immuno-stained wildtype mice with the astrocyte marker GFAP following imaging in a variety of different configurations, and then quantified the immune response of the brain tissue to the corresponding power condition.

Figure S8 shows example stained coronal slices from the different configurations we tested. A negative control condition (Fig. S8a), comprising slices from hemispheres that had undergone surgery for window implantation, but had otherwise not been exposed to laser light, predictably showed no immune response. As a positive control, we imaged mice with ~360 mW of power within a 0.4×0.4×0.5 mm FOV, beyond previously determined safe limits of laser exposure.[24] As shown in Fig. S8b, our positive control condition indeed led to astrocyte activation and an apparent immune response, as expected.

With regards to the acceptable thresholds for the total used laser power, it is worth pointing out that all previous studies of heat-related damage[24,57] have employed significantly smaller cranial windows and smaller FOVs than those used in our experiments. Therefore, it was not immediately evident that previously-established limits for the total power that can be safely used in imaging would apply to our experiments. Accordingly, we conducted further imaging with three additional conditions that represented the most extreme conditions regarding both total power and intensity employed during our imaging experiments: 250 mW delivered within a 3×5×0.5 mm volume (S8c, equivalent to the experiments shown in Fig. 2), 450 mW delivered within a 3×5×0.5 mm volume (Fig. S8d, equivalent to the highest average power used for all experiments), and 250 mW delivered within a 0.6×0.6×0.5 mm volume (Fig. S8e, equivalent to the highest intensity used for all experiments). In all cases, no immune response was observed, indicating that all power levels and intensities used for the experiments shown in this manuscript are within the safe limits for imaging. Figure S8f further shows quantification of the immune response in all above conditions in addition to the positive and negative controls, for $N$ = 3 replicants. Only the positive control condition indicates immunohistochemical response.

While these results demonstrate that the power levels used for our experiments were indeed within safe limits, we also further investigated the relationship between power, FOV, and cranial window size on brain-heating as a means to estimate the safe limit for laser exposure for the large

FOV recordings that our method enables. To that end, we used a finite difference simulation to model heat conduction, metabolic heating, and cooling through perfusion of blood within the brain to estimate the temperature during imaging experiments. We adapted the model from Stujenske *et al.*,[56] including modifications accounting for the intensity pattern generated by laser scanning (l = 960 nm), the geometry and thermal diffusivity of the cranial window, and heat dissipation through the immersion water (assuming a constant temperature of 25° C at a distance 1.5 mm above the cranial window).[57] Additional model parameters are specified in the methods section.

Figure S8g shows an example simulation for when a small FOV (0.4 mm) within a cranial window with an 8 mm diameter is imaged using 450 mW of optical power, a condition that corresponds to the maximum throughput of our excitation laser out of the objective. In the steady state condition, this optical power leads to a brain temperature of ~43 °C, corresponding to an increase of ~6 °C above the core temperature of the animal and likely irreversible adverse effects to the tissue. Additionally, we note that the majority of heating occurs below the focal plane of the microscope (denoted by the magenta line in the figure), in agreement with our immunostaining results which show increased response for tissue below the focal plane for the positive control condition (~600 μm below the surface of the brain, Fig. S8b). In comparison, Fig. S8h shows a second simulation for a 4 mm FOV (8 mm cranial window diameter, approximately equivalent to the experiments in Fig. 2), also illuminated with 450 mW of optical power. In this condition, the observed increase in brain temperature is only by ~1 °C, which we expect, given the results of the condition 2 experiments (Fig. S8d), is within the brain-safe exposure limit.

Using this model, we simulated the maximum temperature increase in the brain as a function of imaging power for a variety of different FOV sizes (Fig. S8i, cranial window diameter = 8 mm). Considering a tolerable brain heating limit of ~3 °C (based on the upper-bound of ethological, activity-induced hyperthermic temperature fluctuations in the mouse brain[57,58]), we can see that the maximum safe limit for laser power ranges from 350-650 mW depending on the size of the FOV, indicating that heating is – at least partially – a function of the imaging intensity. However, given that this safe limit increases by only a factor of ~2 for a 16-fold increase in area, we conclude that the relationship between temperature increase and intensity is sub-linear.

It is also worth noting that the simulations in Fig. S8i show a linear relationship between power and temperature increase for each FOV condition (consistent with previous reports[57]), but only after reaching a certain threshold in power. Looking at Fig. S8h, we can see the source of this threshold: For low power conditions, heat dissipation through the window actually decreases the steady state temperature of the brain below the core temperature of the animal. The effects of brain cooling due to window implantation have been described previously in the literature,[59] with some authors even advocating for heating microscope immersion water to 37 °C in low power imaging experiments in order to maintain core temperature throughout the brain. For our high-power imaging experiments, window-related cooling counteracts laser-induced heating, and allows for increase in the optical power that can be used prior to the onset of the limit for heat-induced tissue response. Additionally, we expect that the initial temperature of the brain in our rodents is cooler in comparison to animals imaged with conventional, smaller FOV microscopes due to increased heat dissipation through the large cranial windows (8 mm diameter) employed in our experiments. This cooler initial condition creates a buffer such that more laser-induced temperature increase in the brain is required to bring the tissue above the core temperature of the animal; thus, rodents with larger cranial windows can likely tolerate higher laser powers without causing adverse effects to the tissue. To explore the relationship between window size and brain temperature, we conducted further simulations for a variety of FOVs, for a fixed laser power of 250 mW. Our results in Fig. S8j show that, increasing the window size to 8 mm can offset heating of the brain during imaging by as much as 1 °C relative to typical 3 mm windows.


58. E. A Kiyatkin, Brain hyperthermia as physiolical and pathological phenomena, *Brain Res. Rev.* ***50***, 27–56 (2005).
59. A. S. Kalmbach, J. Waters, Brain surface temperature under a craniotomy, *J. Neurophisiol.* ***108***, 3138–3146 (2012).
